# MS2Prop: A machine learning model that directly generates *de novo* predictions of drug-likeness of natural products from unannotated MS/MS spectra

**DOI:** 10.1101/2022.10.09.511482

**Authors:** Gennady Voronov, Rose Lightheart, Abe Frandsen, Brian Bargh, Sarah E. Haynes, Elizabeth Spencer, Katherine E. Schoenhardt, Christina Davidson, Andre Schaum, Venkat R. Macherla, Erik DeBloois, David Healey, Tobias Kind, Pieter Dorrestein, Viswa Colluru, Thomas Butler, Marvin S. Yu

## Abstract

Mass spectrometry (MS) is a fundamental analytical tool for the study of complex molecular mixtures and in natural products drug discovery and metabolomics specifically, due to its high sensitivity, specificity, and throughput. A major challenge, however, is the lack of structurally annotated mass spectra for these applications. This deficiency is particularly acute for analyses conducted on extracts or fractions that are largely chemically undefined. This work describes the use of mass spectral data in a fundamentally different manner than structure determination; to predict properties or activities of structurally unknown compounds without the need for defined or deduced chemical structure using a machine learning (ML) model, MS2Prop. The model’s predictive accuracy and scalability is benchmarked against commonly used methods and its performance demonstrated in a natural products drug discovery setting. A new cheminformatic subdiscipline, quantitative spectra-activity relationships (QSpAR), using spectra rather than chemical structure as input, is proposed to describe this approach and to distinguish it from structure based quantitative methods.

## INTRODUCTION

Mass spectrometry (MS) has emerged as the premier analytical tool in a number of disciplines due to its unparalleled combination of high throughput, sensitivity, and mass accuracy.^1–3^ These attributes have led to the generation of large datasets of high information density that are ideally suited for the interrogation of complex biologically derived samples such as those encountered in proteomics and metabolomics. The generation of these large datasets has been complemented by advances in machine learning (ML) leading to the increased use of MS data as an input format for many processes within drug discovery.^4,5^ A significant limitation, particularly in metabolomics and natural products drug discovery, however, is the relative paucity of structurally annotated MS spectra. Publicly available databases of natural products and metabolites such as COCONUT^6^ and the Metabolomics Workbench Metabolite Database^7^ contain on the order hundreds of thousands of compounds, while structurally annotated MS spectra available through GNPS,^8^ MoNa,^9^ NIST Mass Spectral Libraries,^10^ or MassBank^11^ measure in the tens of thousands, each representing a fraction of the billions of estimated natural metabolites.^12^ The vast majority of metabolites, therefore, remain structurally opaque such that the MS analysis of metabolomic samples reveals large numbers of metabolites, but most of them of undetermined structure, properties, and function. This has hampered natural products drug discovery as traditional bioactivity guided fractionation approaches often result in the isolation and identification of molecular non-starters from a drug discovery and development standpoint. Methods by which bioactive natural products can be prioritized for isolation and identification based on drug or lead-like properties using MS spectra without requiring structurally annotated spectra would be highly desirable to improve natural products drug discovery output and efficiency.

The prediction of structure or properties from MS spectra, however, has relied primarily on either matching spectrum to reference spectra from molecules of known structure or molecular networking^13^ unknown compounds to those reference spectra to provide structural clues. It is only through association with known annotated spectra that structure or property predictions for non-annotated spectra can be made through quantitative structure-activity or property relationships (QSAR/QSPR). These methods mathematically model chemical properties and biological activity to structural morphology based on the principle that structural similarity leads to similar activity and properties. The advent of machine learning, deep learning, and other advanced computational methodologies has accelerated the use and application of QSAR^14^ in a number of fields including drug discovery^,15,16^ but the relative paucity of annotated reference spectra limits the ability to examine undefined chemical space.

QSAR and QSPR, by definition, require a chemical structure as the data input in the form of a molecular representation, generally a SMILES string, a molecular fingerprint, or a molecular graph. These molecular representations are used to generate molecular descriptors that are correlated to the desired chemical property or activity.^17^ The dependency on chemical structure presents its own issues, since molecular representations are imperfect, either in their ability to depict certain molecular configurations, such as tautomers or aromaticity, in an unambiguous manner.^18^ Natural products have been cited as particularly difficult for molecular representations due to their large molecular weight range, multitude of stereocenters, and high fraction sp3 content.^19^ These challenges with molecular representations have been shown to substantially impact QSAR predictions.^20^

In addition, a recent editorial highlighted that a primary obstacle to applying ML to predict chemistry has been the lack of good training sets.^21^ While speaking specifically to the application of ML to retrosynthesis, the following can apply generally, “To make accurate chemical predictions, an AI system needs sufficient knowledge of the specific chemical structures”. The acquisition of good training sets with enough specific chemical structures can be a primary challenge for many QSAR exercises.

Approaches to chemical property or biological activity modeling that require neither starting from a known chemical structure nor comparison to reference spectra have been sporadic and limited in scope. A simple early example, analogous to initial regression analysis QSAR approaches, is the use of liquid chromatography retention data to predict log p.^22^ Recently, however, there have been several interesting machine learning approaches to the prediction of chemical properties of analytes of unknown composition, primarily within the environmental sciences. Franklin and colleagues^23^ analyzed atmospheric samples by GC-ESI-MS and used both chromatographic and spectral data to develop models based on random forest methodology to predict properties of unknown compounds such as carbon:oxygen ratio, average carbon oxidation state, and vapor pressure. Zushi, meanwhile also used GC-MS data and a decision tree-based learning system, XGBoost, to develop predictions of various chemical properties directly from spectra, including logK_o–w_, molecular weight, and water solubility. Zushi^24^ went further and predicted rodent toxicity directly from the GC-MS data, but with marginal success. While these efforts were noticeably constrained to compounds amenable to gas chromatography and primarily to datasets that do not contain large amounts of co-eluting or isomeric compounds since only MS1 and retention time data is used, Boelrijk *et al*.^25^ used MS^2^ fragmentation data to predict retention indices for unknown compounds using another decision tree regression model, CatBoost. Peets *et al*. ^26^ also used MS^2^ spectra to build ML-based predictive models of environmental toxicities and concentrations of unknown compounds in water samples, although there was still some dependency on spectral comparisons and structural representations such as SMILES strings. Together these reports provide a foundation for further efforts in using non-structure based methods for the development of predictive property and activity models that are decoupled from molecular structure and allow the exploration of structurally undefined chemical space.

### The key concept in using MS/MS data as an input format for activity relationships, is that spectra are, in essence, computer-ready molecular representations

MS/MS spectra are the result of fragmentations based on atomic composition, disposition, and arrangement such that every spectrum, to some degree, has information embedded within it similar to that contained within a molecular representation such as a SMILES string or chemical graph. David *et al* defined a chemical representation as “any encoding of a chemical compound”.^27^ MS/MS spectra clearly fit this straightforward definition. This relatively simple, but critical awareness provides the basis for a new subdiscipline, dubbed quantitative spectra-activity relationship (QSpAR), uniquely differentiated from QSAR/QSPR by freeing the analysis from chemical structure and therefore allowing for the interrogation of large datasets that contain little or no prior structural annotation.

The expansion of the QSpAR concept to natural products drug discovery using LC-MS/MS data and large language models is presented herein as MS2Prop, a ML model by which ten chemical properties used in drug discovery are predicted without the requirement for an intermediate structure or reference spectra. The architecture of MS2Prop is detailed, property predictions both in and out of the training set reported and validated, and performance benchmarked to structure-dependent models. Lastly, the performance of MS2Prop is evaluated in a real-world scenario by predicting properties of compounds contained in a plant-derived complex mixture.

## RESULTS AND DISCUSSION

### MS2Prop model architecture and training

MS2Prop predicts 10 numerical chemical properties; atomic logP,^28^ number of hydrogen bond acceptors, number of hydrogen bond donors, polar surface area, number of rotatable bonds, number of aromatic rings, number of aliphatic rings, fraction of sp3 carbons, quantitative estimate of drug likeness,^29^ and synthetic accessibility^30^. These properties were selected for their relevance to drug discovery and medicinal chemistry and are easily computed within RDKit^31^ from chemical structures which will allow for benchmarking MS2Prop’s performance against other methods.

Conceptually, MS2Prop maps MS/MS spectra to chemical properties by aggregating information across the fragment peaks to create a latent vector representation of the input spectrum and using this vector to predict the target properties. The model is built around a deep neural network architecture consisting of three main stages (**Fig. 1**). First, the input spectrum – represented as a sequence of *m*/*z*-intensity pairs along with the precursor *m*/*z* – is mapped to a sequence of embeddings, i.e., dense real-valued vectors. Next, the sequence of embeddings is passed through a transformer.^32^ A critical component of the transformer architecture is that each layer of the network has an output for each element of the input sequence and can weight those outputs dynamically in the most relevant way for learning. While transformers have largely been associated with natural language processing tasks, they have also seen application in a large number of diverse applications^33,34^ and are a natural choice for adaptive aggregation of information across the peaks of MS/MS spectra. The final layer of the transformer outputs a sequence of vectors with MS2Prop only retaining the first output vector, which can be considered a latent embedding for the entire MS/MS spectrum. Finally, the spectrum embedding is passed to a prediction head, which is a feed forward neural network that outputs the predicted chemical properties. Because this final prediction module computes all properties, only a single inference call is needed. Importantly, at no time does the model rely on comparison to annotated spectra, reference any external database, nor deduce structures to calculate the properties. The generated predictions are, therefore, completely *de novo* and rely only on MS^2^ spectral data.

**Figure 1:**
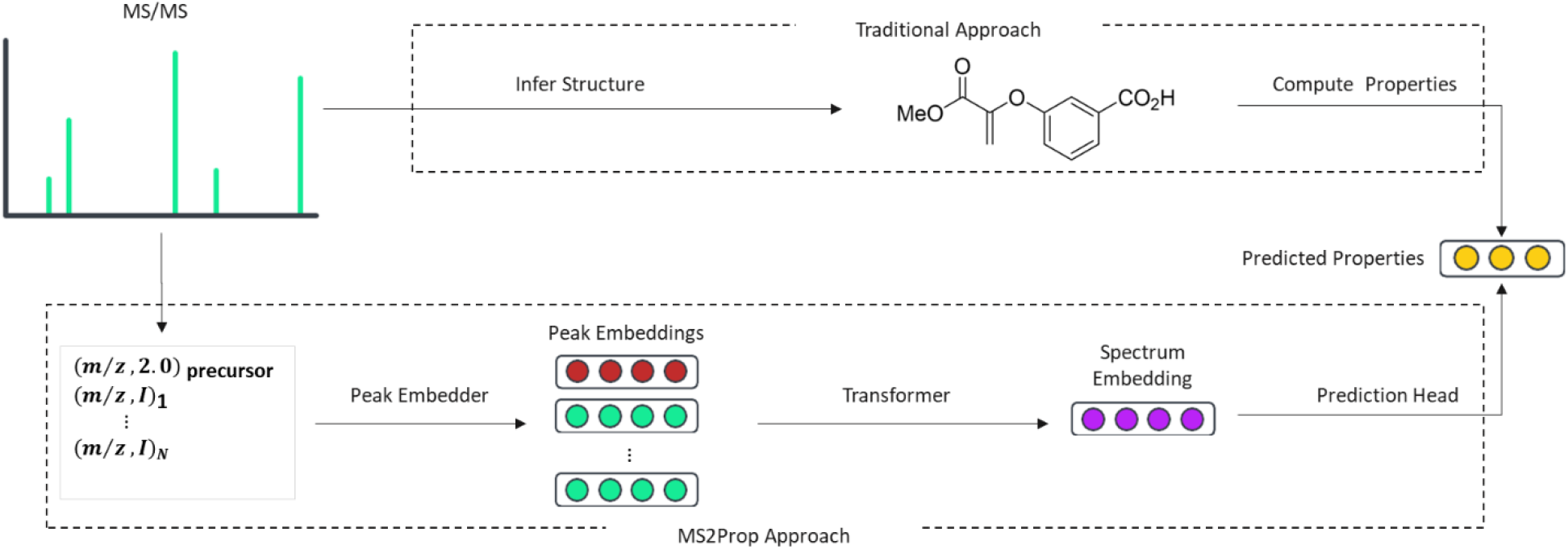
Overview of MS2Prop architecture for property prediction compared to structure-based approaches.

The entire MS2Prop model was trained by minimizing the mean squared error between the predicted and true properties over a labeled training dataset of annotated spectra from a number of publicly available datasets.^8,10,11,35–43^ Spectra collected in both positive and negative ion-mode were included, in addition to a variety of instrument types, collision energies, and other instrument parameters..

### MS2Prop performance and benchmarking

Traditionally, machine learning systems are often validated using a single test set that is simply a random sample from the same pool of data used in training. However, work on molecular property prediction from chemical structures has shown that such test sets over-estimate generalization performance because the data is too similar to the training data compared to realistic use cases.^44^ This phenomenon is common within machine learning in complex domains, and similar observations have been made in natural language processing.^45^ To address this, three different test sets of MS^2^ spectra were deployed with each an increasingly stringent test of performance to accurately estimate the generalization performance of MS2Prop. The first test set (998 spectra corresponding to 941 distinct compounds) consists of MS^2^ spectra of compounds contained within the MS2Prop training set, but the spectra themselves were not in the training set. This “known” test set represents the easiest generalization setting. The second “novel” test set consists of 17,270 spectra that correspond to 1,002 compounds that were not contained in the MS2Prop training set and is a step up in difficulty from the known test set. The final, and most challenging, test set is the CASMI 2022^46^ set which were generated in an independent experimental setting, and whose 499 spectra corresponding to distinct molecules were chosen because they were not in spectral reference libraries. A key motivation for using the CASMI2022 dataset is that it is assured to be distinct both in structure and experiment from any training data used for MS2Prop or any of the comparator methods to be used for benchmarking. In all cases, predicted structures are contained in public databases such as PubChem that were available to CSI:FingerID at inference time, but which were not accessed by MS2Prop.

MS2Prop was fed a single MS^2^ spectrum for each molecule within the three test sets and the model provided the values for each of the selected properties. MS2Prop’s performance for each property and test set as measured by *R*^*2*^ when compared with the RDKit calculated values for each compound in a test set is shown in **Table 1**. For the known test set, MS2Prop can reproduce the properties of the compounds from their MS^2^ spectra nearly perfectly (*R*^2^ = 0.96). For the novel test set, MS2Prop performance aggregated across 10 properties is *R*^2^ = 0.*8*0 and for the CASMI test set it was *R*^2^ = 0.*79*. MS2Prop can, therefore, be used to predict properties for compounds based solely on MS^2^ spectra with no structural reference or comparison required with a reasonable degree of accuracy overall.

**Table 1:** MS2Prop Performance (*R*^2^) for Predicted Chemical Properties by Individual Test Sets.

| Property | Known test set | Novel test set | CASMI 2022 |
| --- | --- | --- | --- |
| atomic logP (AlogP) | 0.95 | 0.63 | 0.61 |
| number of hydrogen bond acceptors (HBA) | 0.98 | 0.94 | 0.93 |
| number of hydrogen bond donors (HBD) | 0.96 | 0.75 | 0.85 |
| polar surface area (TPSA) | 0.98 | 0.92 | 0.91 |
| number of rotatable bonds (# Rot. Bonds) | 0.96 | 0.85 | 0.67 |
| number of aromatic rings (# Arom. Rings) | 0.97 | 0.67 | 0.75 |
| number of aliphatic rings (# Aliph. Rings) | 0.96 | 0.81 | 0.80 |
| fraction of sp3 carbons (Fsp3) | 0.97 | 0.84 | 0.79 |
| quantitative estimate of drug-likeness (QED) | 0.94 | 0.76 | 0.78 |
| synthetic accessibility (SA) | 0.96 | 0.82 | 0.77 |
| all properties aggregated | 0.96 | 0.80 | 0.79 |

In terms of individual properties, MS2Prop performed best for number of hydrogen bond acceptors, polar surface area, and number of aliphatic rings with an *R*^2^ ≥ 0.*80* on both the novel and CASMI dataset. The most challenging property for MS2Prop is atomic logP, however MS2Prop still had a respectable *R*^2^ > 0.61 on the two more challenging datasets. An alternative approach to property predictions from MS^2^ spectra, and one that is most often used currently, is to first predict molecular structure from the MS^2^ spectra and then calculate properties from these predicted structures.^47,48^

In order to benchmark MS2Prop, two standard approaches for structure identification from MS^2^ data; a modified cosine similarity spectral lookup^49^ and CSI:FingerID^50^ were evaluated and properties calculated from the predicted structures. The modified cosine similarity spectral lookup was used as an initial benchmark for MS2Prop, since a direct comparison is possible between the two methods with all three datasets (**Fig. 2**). As shown, MS2Prop performed significantly better across all datasets and properties (**Fig. 2b-k**). For some properties, such as the number of hydrogen bond acceptors (**Fig. 2c**), both methods have performed well, while others such as atomic *log*P, proved more challenging, but even in that case the stronger performance of MS2Prop was clearly evident over modified cosine similarity.

**Figure 2:**
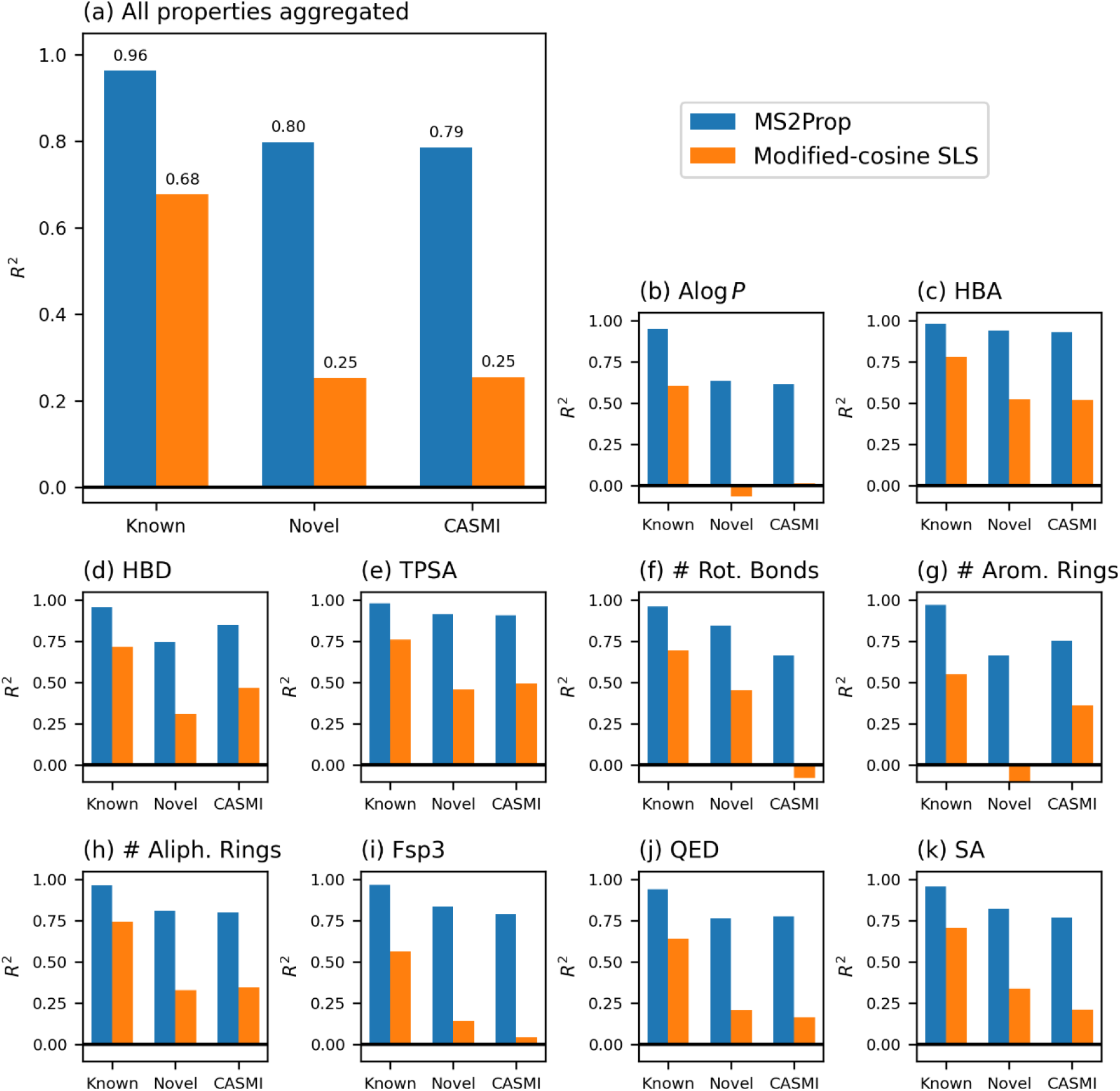
Model performance as measured by R2 across all properties (higher is better). R2 results are aggregated across properties in (a) and shown individually in (b)-(k). Performance is reported across three test datasets: known, novel, and CASMI 22 and two models: MS2Prop and cosine similarity spectral lookup.

Direct comparison with CSI:Finger ID is not possible for the known and novel datasets, since the training set used for CSI:Finger ID is unknown but is possible using the CASMI2022 dataset which is naive for both methods. In this most challenging dataset MS2Prop outperformed CSI:FingerID against all 10 properties and both MS2Prop and CSI:Finger ID outperformed modified cosine similarity by a wide margin (**Fig. 3**). Once again variation in individual properties was observed with atomic log p and number of rotatable bonds being particularly difficult for CSI:Finger ID.

**Figure 3:**
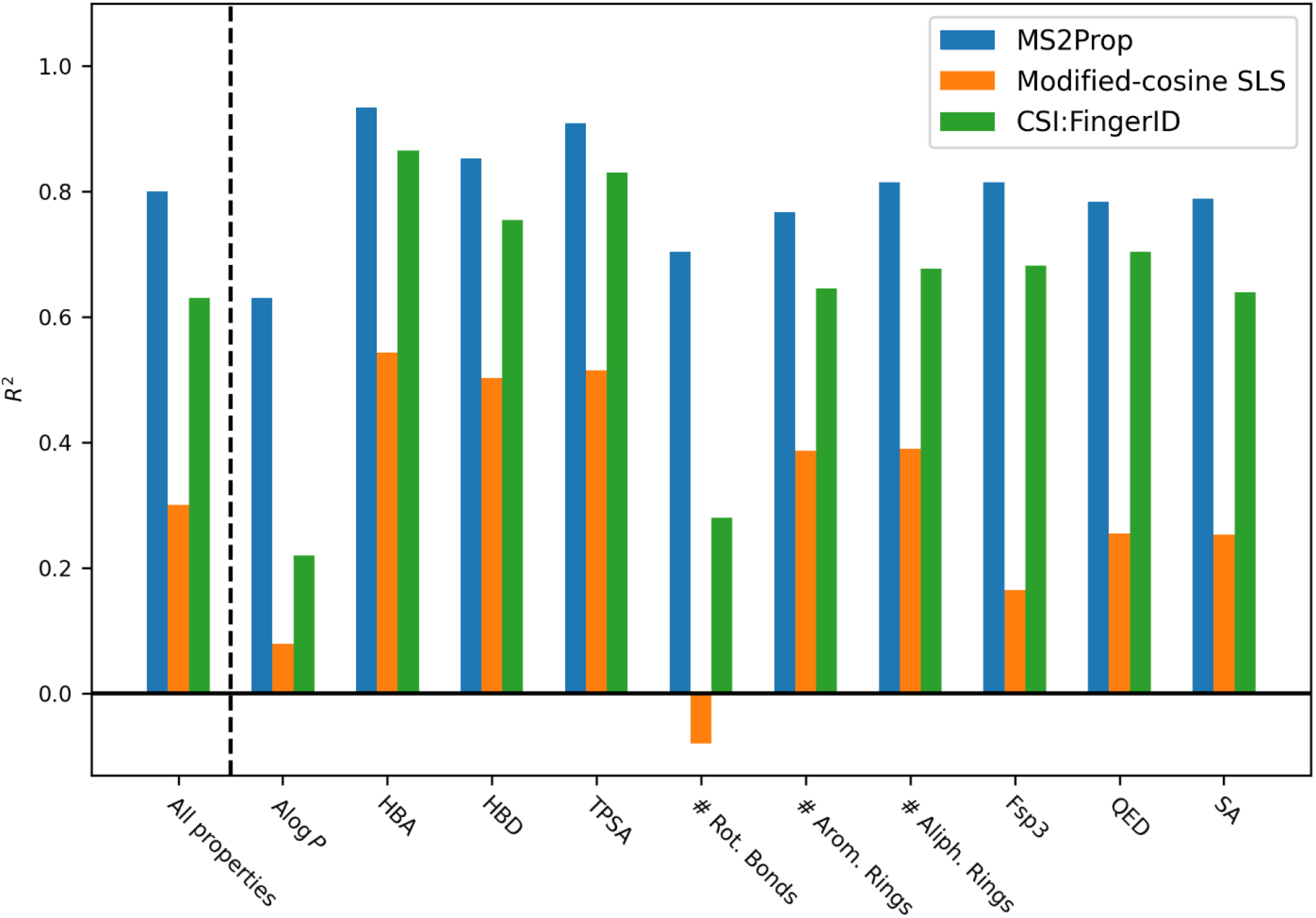
Performance comparison across various approaches on CASMI dataset. MS2Prop, Modified Cosine similarity spectral lookup, and CSI:FingerID model performance as measured by R2 across all properties (higher is better) reported on the subset of CASMI 22 spectra for which CSI:FingerID was able to produce structure predictions.

Two of the ten properties that should be highlighted are Quantitative Estimate of Drug-likeness (QED) and Synthetic Accessibility (SA), that are both multi-parameter calculations generally requiring that a number of descriptors or structural characteristics first be calculated from a structure then be combined to provide a single value. The MS2Prop model, however, predicts these values without any intermediate parameters required. Synthetic accessibility, in particular, is interesting, since a major component involves breaking a structure representation down into fragments through the use of molecular representations which MS/MS fragmentation does physically to a compound. This demonstrates that QSpAR approaches can be used to directly derive values that are usually an ensemble of various properties or descriptors. It is envisioned, therefore, that the model can be trained on chemical and biological properties, such as blood brain penetrance or biological activities, which would be of interest to the drug discovery community. Coupled with the ability to quickly generate large training sets without the necessity of definitive structure determination would in theory accelerate the building of QspAR models compared to traditional QSAR/QSPR approaches.

While predictive performance between CSI:FingerID and MS2Prop consistently favors the latter, this is only part of the total performance advantage of MS2Prop. First, CSI:FingerID fails to deliver any output at all in certain instances as it did not provide a prediction for 36 of the 499 spectra(7%) in the CASMI2022 dataset. The data in Fig. 3 excludes these null predictions and only includes those instances when the model produced an answer. Second, the difference in the overall computational time needed for CSI:FingerID vs MS2Prop was substantial (**Table 2**). On a per prediction basis, MS2Prop averaged 36.8 milliseconds while CSI:Finger ID averaged 27.3 seconds for those instances when an output was delivered. Extrapolation of the timings in Table 2, MS2Prop would take 10 hours to generate property predictions for a million unannotated spectra while the same task would take CSI:FingerID over 8.5 days. Across both novel and known compounds MS2Prop’s performance is substantially higher, both in terms of predictive performance and scalability than either modified cosine similarity score or CSI:Finger ID.

**Table 2:** Time to predict properties on all 499 spectra in CASMI2022 test set.

| Model | Elapsed time |
| --- | --- |
| MS2Prop | 18.35 seconds |
| CSI:FingerID | 368 seconds |
| Modified cosine SLS | 7.36 hours |

### MS2Prop performance using spectra from plant extracts

To further evaluate MS2Prop’s performance, we used MS/MS spectra obtained directly from a complex mixture, rather than spectra obtain on pure compounds as in the test cases. An ethanolic extract of *Withania somnifera*, a well-studied plant with an array of bioactivities attributed to it,^51^ was used and 26 natural products were spiked into the extract prior to untargeted LC-MS/MS analysis of the extract. In addition to these 26 known compounds that were spiked in, nine known compounds from the *W. somnifera* extract were isolated after LC-MS/MS profiling and their structures confirmed via NMR. The 26 exogenous and 9 endogenous compounds were separately profiled on the same MS instrument (i.e., not with the plant extract background), and this information was used to locate the MS features for each of these 35 compounds within the LC-MS/MS run of the complex mixture. The MS2Prop predictions derived for the 35 compounds, the “extract” dataset, based solely on their MS^2^ spectra collected during the untargeted profiling of the complex mixture were then compared with calculated values for the ten properties based on known structure (**Fig. 4**). Importantly, the total MS^2^ signal counts for all 26 exogenous compounds were well below that of the highest isolated compound (Supplemental Information, Table S1) indicating that the experiment accurately reflects an untargeted LC-MS/MS profiling run containing compounds in varied abundance.

**Figure 4:**
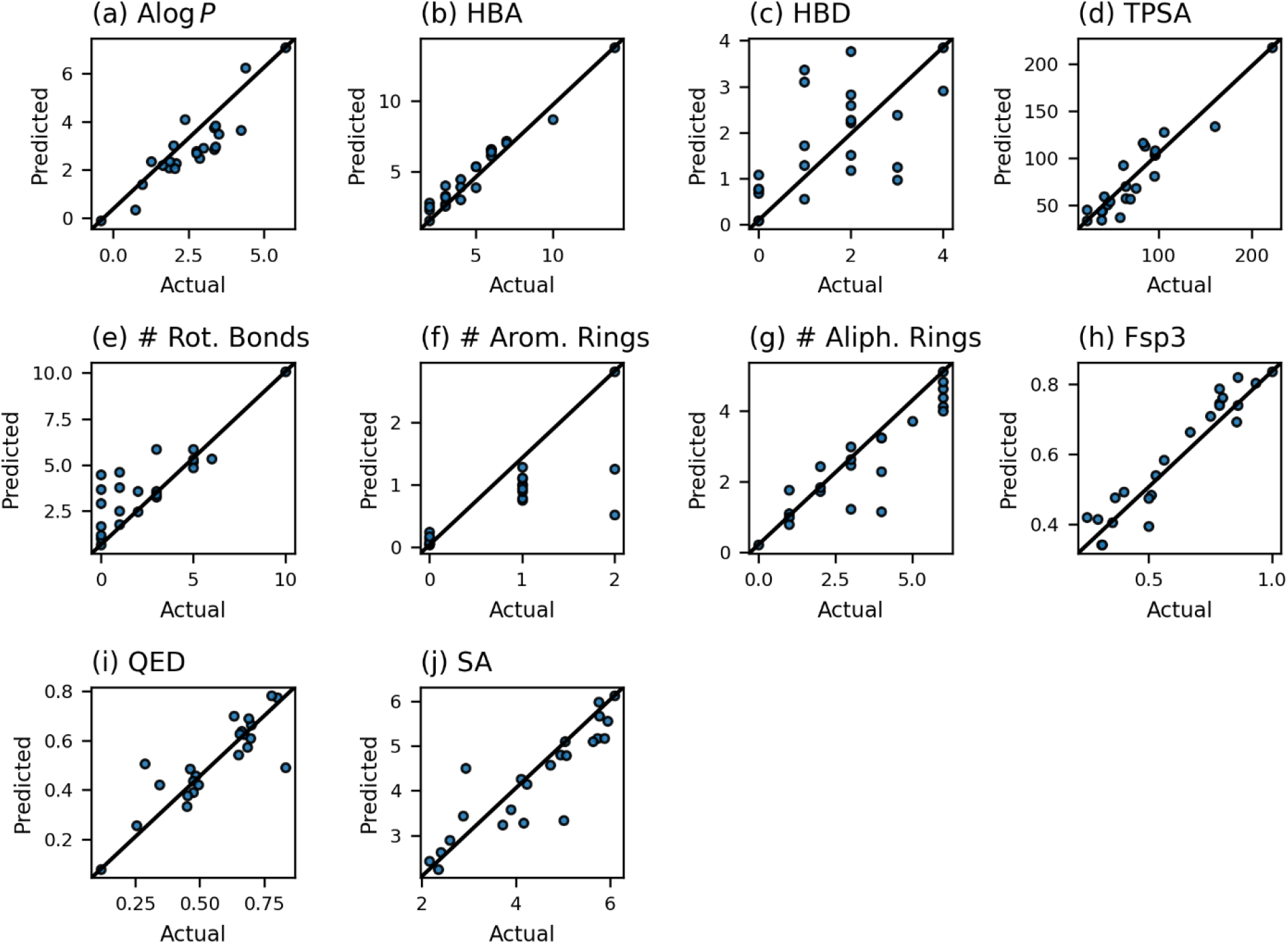
Predicted properties from MS/MS spectra from a complex mixture vs properties calculated from known structures. These are predicted on [M+H]^+^ MS2s obtained on both the exogenous compounds added to the extract and endogenous compound isolated from the extract.

The performance of MS2Prop using MS^2^ spectra collected from an extract profiling run was then compared to using MS^2^ spectra from pure compounds in the CASMI 2022 dataset. While *R*^2^ was used to compare the performance of MS2Prop, CSI:Finger ID, and modified cosine similarity on the three prior datasets, to compare performance between the extract and the CASMI 2022 datasets the ratio of mean absolute error (MAE) for each property was used (**Fig.5**) since *R*^2^ is not an appropriate metric across distinct datasets. As seen, there was a slight decrease in performance in the experimental setting compared to the test setting with 5 of the properties showing statistically significant (*p*<0.05) increases in mean absolute error. The absolute value of these errors, however, is mostly within a reasonable range for decision making purposes in terms of identifying metabolites with drug-like properties. MS2Prop’s predictive ability is, therefore, considered translatable to the demands of natural products drug discovery.

**Figure 5:**
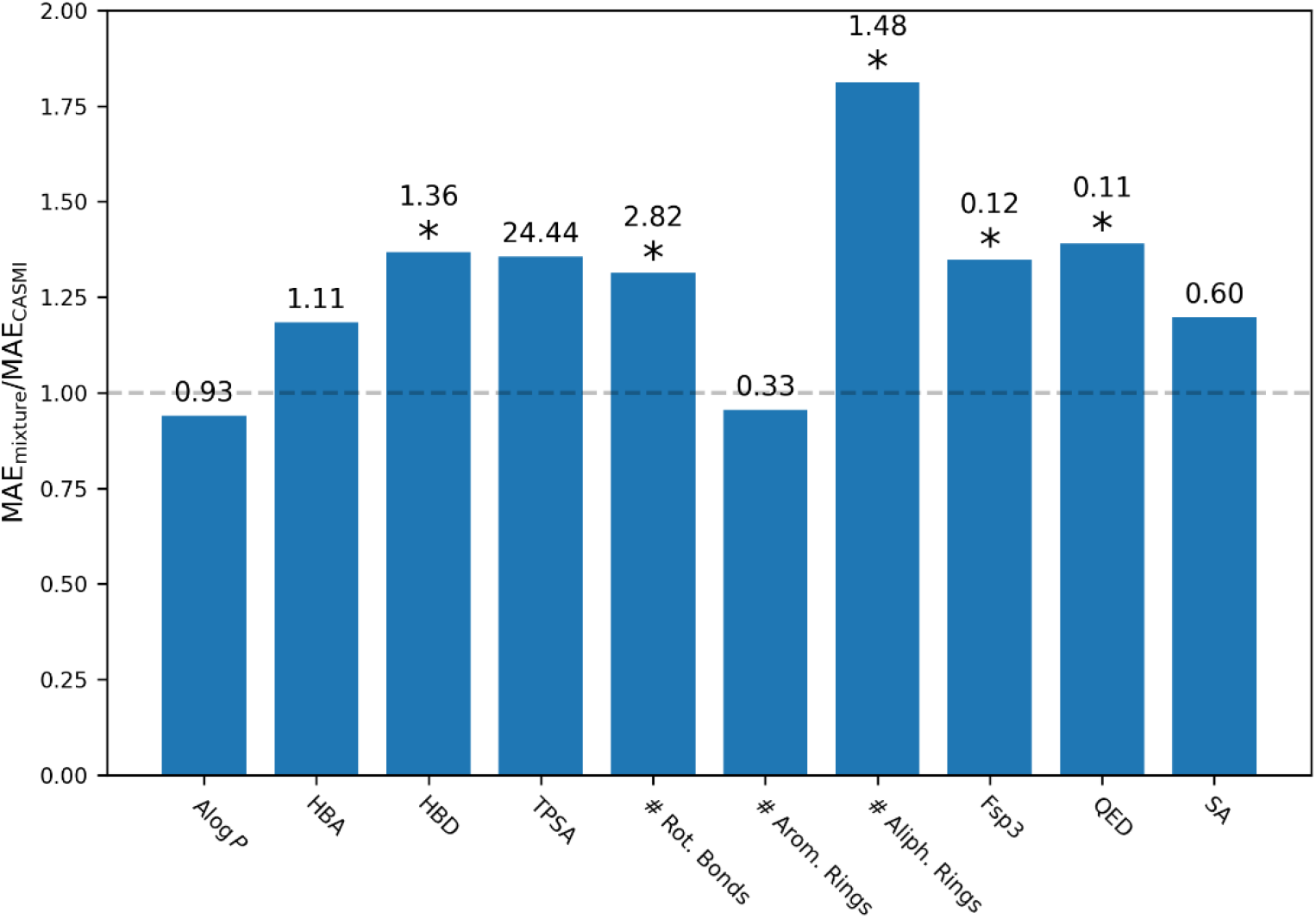
Relative MAE of MS2Prop predictions on *W. somnifera* extract dataset vs. CASMI 2022 dataset. The value above each bar is the MAE for the extract dataset. *Statistically significant difference between extract MAE and CASMI 2022 datasets (one-sided t-test p-value less than 0.05).

The approach of directly predicting properties using MS^2^ spectra instead of predicting the compounds or fingerprints as intermediate objects has two principal advantages. The first is that it avoids complex, sometimes non-monotonic, propagation of errors that arise in multi-stage machine learning systems.^52^ The second principal advantage is that fingerprints or compound structures are high dimensional objects that are far more complex than the scalar properties predicted by this model. Avoiding the need to predict these complex intermediate objects, only to go back to scalar properties, increases the tractability of the learning problem, and likely plays a large role in the high performance of MS2Prop, both in terms of predictive accuracy and scalability. These factors in conjunction with the use of modern training and transformer architecture pioneered in large language models^32,53^ likely drive MS2Prop’s strong performance and generalization to unseen structures.

Not surprisingly, since CSI:Finger ID has primarily been used as a structure determination algorithm, the first step is to transform MS^2^ fragment spectra into a chemical formula fragment tree which is then processed over several steps into molecular fingerprints. MS2Prop, on the other hand, treats the MS^2^ spectra itself as a molecular representation thereby eliminating the fragment tree formation and processing steps altogether. By using large language models, a spectrum is akin to a sentence where words (individual peaks) are deduced through context without requiring translation (fragment assignment). The QSpAR approach, therefore, expands the use of mass spectrometry data beyond its two most prevalent uses; a) structure determination and b) analyte detection and quantification and into direct chemical property and bioactivity interrogation.

Even with MS2Prop’s robust performance, there remain opportunities for improvement that remain unexplored. The current implementation incorporates molecular formula information only indirectly using the precursor mass as a feature. Modern tandem MS analysis tools can identify molecular formulas with 90+% accuracy in many contexts.^54–56^ Additionally, the annotated training data used was highly skewed towards certain adducts such as [M+H]^+^. No efforts have been made to correct this bias in the training of MS2Prop and hence it’s performance on other adducts suffers.

The transformer architecture used here, however, is well-suited to self-supervised learning approaches pioneered in large language models that allow the model to learn from abundant unlabeled mass spectrometry data.^53^ The strong performance of the current model is a baseline for future work and prediction accuracy will improve with further development. The architecture also allows other properties to be readily predicted given an appropriate training set. Without the need for defined structure the acquisition of large datasets using structurally undefined but MS^2^ profiled compound mixtures presents the potential to rapidly produce high quality training sets.

## CONCLUSIONS

The generation of large datasets by LC/MS-MS along with the ability to analyze these sizable datasets in an efficient and scalable manner using advanced machine learning techniques represents a powerful strategy by which to explore novel chemistry and biology within natural products drug discovery. A major limitation has been the requirement to either have structurally annotated reference spectra or to translate spectra to a molecular representation. By regarding MS^2^ spectra and their fragmentation patterns that serve as functional equivalents to SMILES strings or graphs, however, this limitation can be overcome as exemplified by the MS2Prop model that can accurately predict chemical properties without the need for intermediate structure representation.

MS2Prop uses natural language processing to accurately predict drug-like chemical and multi-parameter properties. The model was validated using three different test sets of varying difficulty, benchmarked against structure-based predictive methods, and demonstrated in a real-world experimental setting. MS2Prop proved to be superior in terms of predictive accuracy and scalability with the latter being particularly important to properly leverage the throughput and sensitivity of modern mass spectrometry methods. By decoupling chemical structure from property prediction, molecules of potential interest can be identified from plant and microbial extracts, prior to isolation and structure determination to overcome many of the inefficiencies of traditional bioactivity guided fractionation that often led to non-viable hits from a drug discovery standpoint. MS2Prop is an example of a scalable QSpAR model for natural products drug discovery. It is envisioned that the application of the principles and methods described will inspire other QSpAR efforts and accelerate the development of models to correlate spectra to function for drug discovery, biomarker discovery, metabolomics, and other areas of interest, while simultaneously expanding the application of mass spectrometry beyond structure determination and analyte detection.

## EXPERIMENTAL SECTION

### Data Preparation

The following filtering steps were applied to ensure uniform quality datasets. First, spectra with fewer than 3 decimal places of *m/z* resolution were excluded. Next, spectra with precursor *m/z* greater than 1,000 Daltons were excluded. Similarly, from each spectrum, peaks with *m/z* greater than 1,000 Daltons were excluded. For each spectrum, peaks are sorted by intensity and only the top 512 peaks are retained. Finally, spectra that have fewer than 5 peaks remaining after the previous filtering steps were excluded.

For each spectrum, intensities were normalized to a maximum of 1 and m/z values rounded to the nearest 0.1 Da, similar to Spec2Vec,^55^ so that each peak is represented by a discrete token m/z and a normalized intensity value.

All stereochemistry was stripped from molecular structure labels, a common step taken in MS/MS modeling that allows for better molecule-disjoint splitting. Chemical properties and fingerprints were computed from the cleaned molecules using RDKit.^31^ In total, the resulting labeled data contained approximately 1,250,000 spectra corresponding to approximately 45,000 distinct molecules.

### MS2Prop Architecture

#### Model Input

The MS2Prop model treats an input MS/MS spectrum *S* as a set (**Fig. 6, Eq.1**) comprising a precursor *m*/*z* and *N* fragment peaks at various *m*/*z*’s and intensities *I*’s. Since the precursor *m*/*z* indicates the molecular mass and does not have an experimental intensity value, it is always assigned an intensity of 2.0. Hereafter, each tuple (*m*/*z, I*) within a spectrum is referred to as a *peak*.

**Figure 6:**
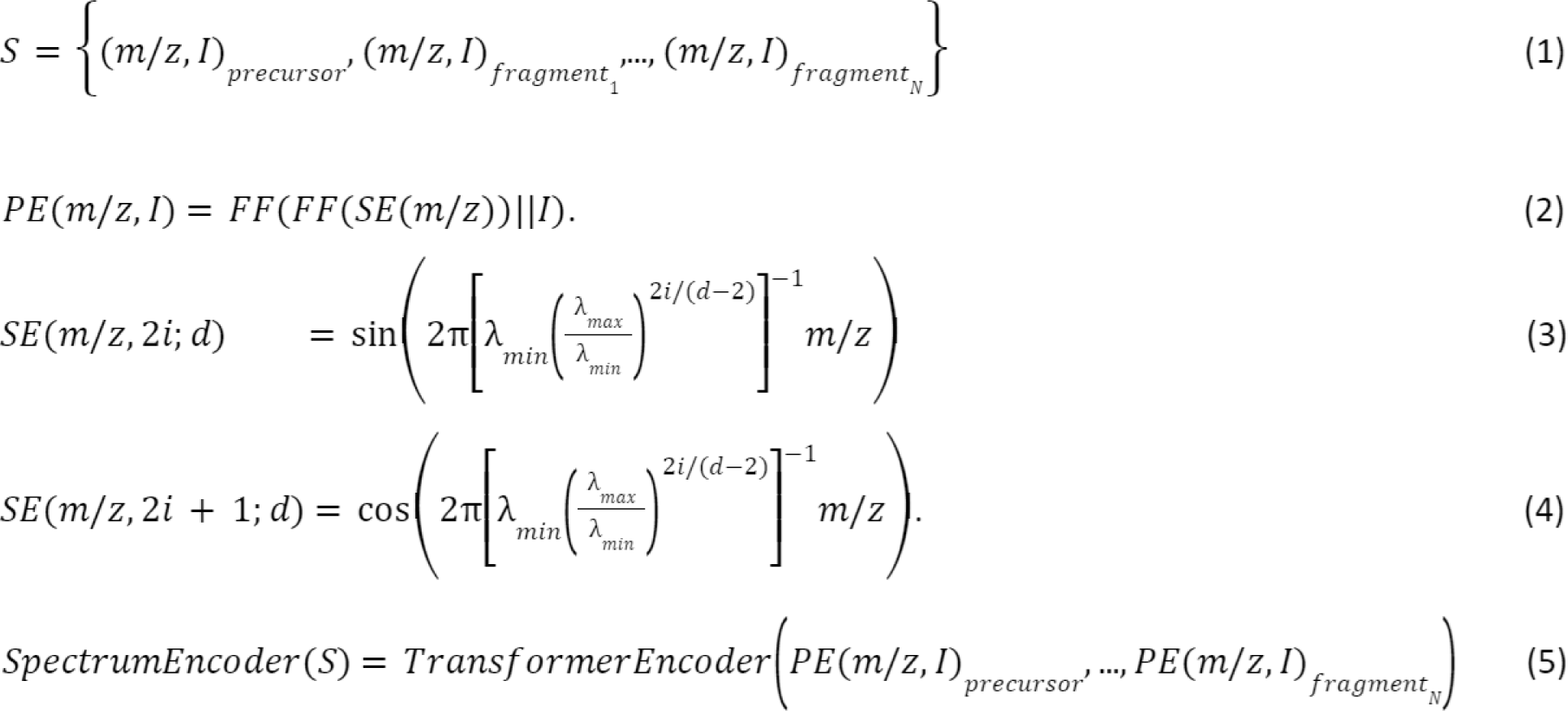
MS2Prop Architecture Equations.

#### Peak Embedding

Given an input MS/MS spectrum *S*, each peak was embedded into a continuous vector space (**Fig. 6, Eq.1**). We employ a sinusoidal embedding,^32,57^ *SE*, to represent high resolution measurements of *m*/*z* as a vector of dimension *d* = 512 (**Fig. 6, Eqs. 3 and 4**).

The frequencies are chosen so that the wavelengths are log-spaced from *λ*_*min*_ = 10^−2.5^Daltons to *λ*_*max*_ = 10^3.3^Daltons, corresponding to the mass scales of interest. Finally, the || operator denotes concatenation and *FF* is the standard feed-forward neural network with a single hidden layer of dimension *d, ReLu* non-linearity, and output dimension *d*. A diagram of *PE* is shown in **Fig. 7**.

**Figure 7:**
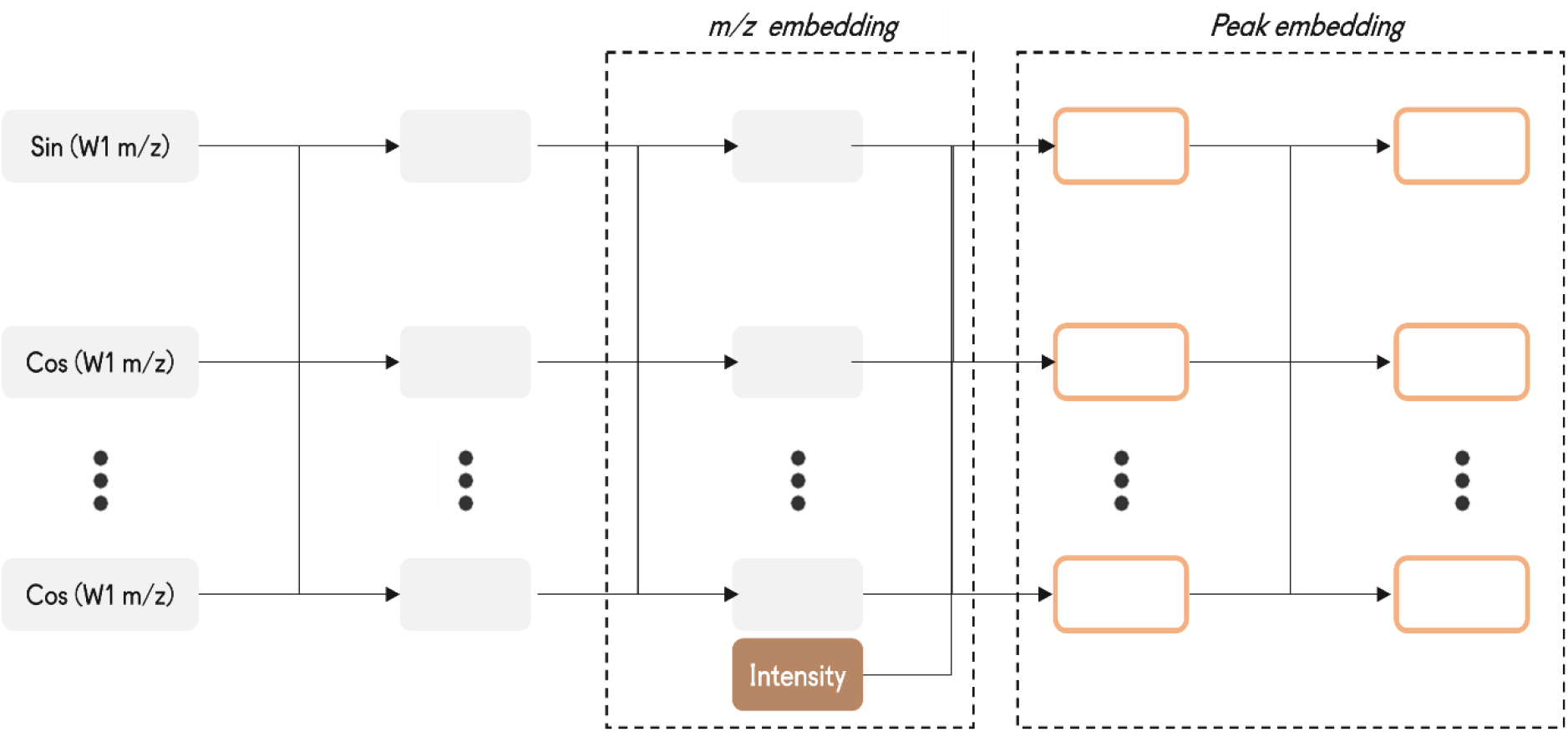
Architecture used to embed MS/MS peaks composed of a m/z and signal intensity.

### Spectrum Transformer Encoder

Transformer encoders without the positional encoding are fully-symmetric functions and hence are ideally suited to model a set of MS/MS fragmentation peaks.^58^ Therefore, MS2Prop passes a sequence of peak embeddings to a transformer encoder (**Fig. 6, Eq. 5**) where, *TransformerEncoder* has embedding dimension *d* and six layers, each with 32 attention heads and an inner hidden dimension of *d*. As mass spectra have no intrinsic ordering, no positional encoding was included.

To get a single embedding vector as an output, the final transformer layer query only attends to the first embedding, corresponding to the position of the precursor *m*/*z*. A diagram of the full model architecture is shown in **Fig. 8**.

**Figure 8:**
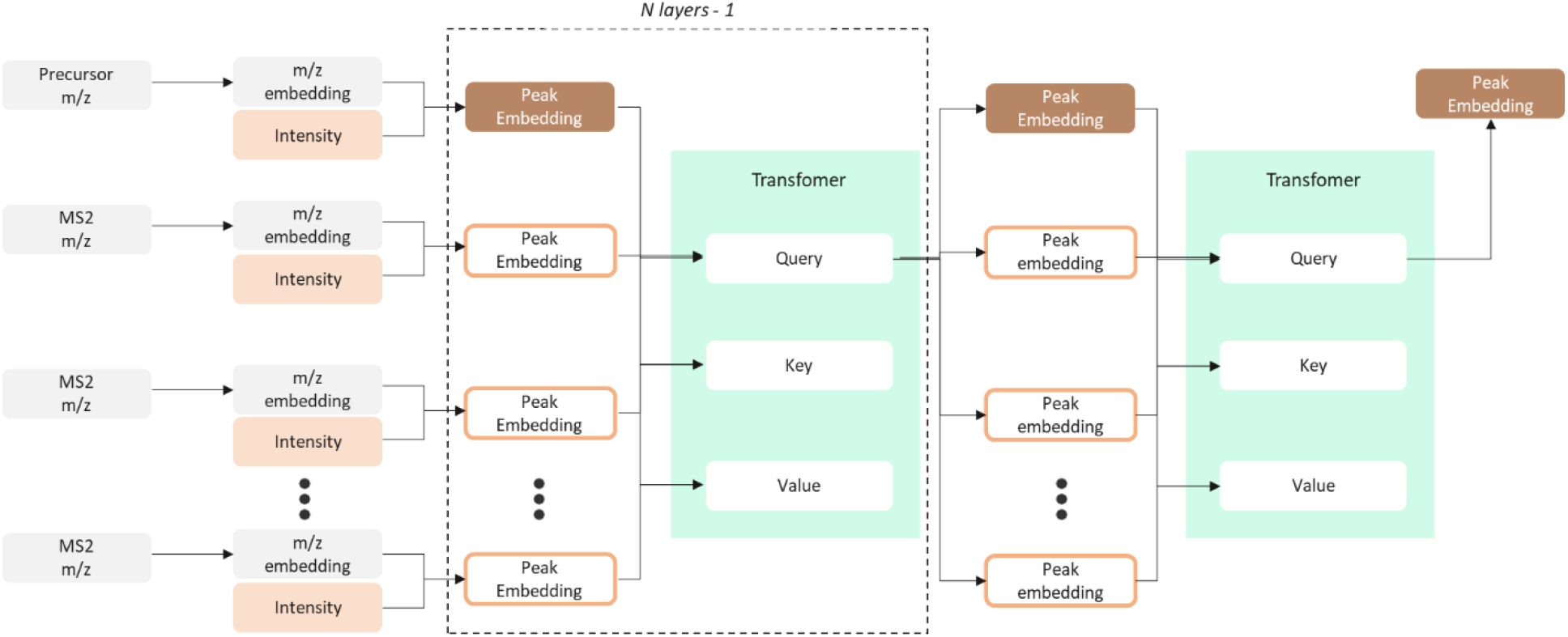
Full model architecture of MS/MS spectrum that are featurized and embedded in a dense vector space.

### Property Prediction Head

After encoding the input spectrum to a single vector representation, MS2Prop passed this to a simple feed-forward neural network with 1 hidden layer of dimension *d* and output dimension 10 to predict the desired properties.

### Model Training

MS2Prop is trained by minimizing the mean square error loss averaged across the 10 properties considered here. During training, the target property values are scaled to zero mean and unit variance. The Adam^59^optimization algorithm with parameters 1=0.9, 2=0.999, and a learning rate of =5.010-5 is used to implement the minimization. Additionally, the following parameters are used, a weight decay^60^ of 0.1, a gradient clipping of 0.5, and a dropout^61^of 0.1. Finally, MS2Prop is trained over 34 epochs with a batch size of 64. MS2Prop is built and trained using PyTorch.^62^

### Modified-Cosine Spectral Library Search

For all modified-cosine results, we used matchms^49,63^ version 0.11.0. Every query spectrum was then matched to an annotated spectra from the MS2Prop training set that had the highest modified cosine spectral similarity (evaluated using default parameters).

### Sirius and CSI:FingerID

For all CSI:FingerID results, we used SIRIUS version 5.6.3 (Bright Giant, GmbH, Jena,Germany). Query data was prepared as an .mgf file. Each spectrum contained a separate entry for MS1 (containing isotope masses and intensities gathered internally through Enveda’s feature calling software) spectra and associated MS/MS spectra. In keeping with all other results, MS/MS spectra were limited to the most intense 256 fragments. Each molecule was represented by at most one spectrum from each available (collision energy, adduct) pair. Both positive and negative ion mode spectra were included. Spectra were evaluated separately, not merged across ion modes, collision energies, or adducts. No timeout was used.

SIRIUS was set to use DB formulas from the Bio Database only, with all possible default ionizations. The instrument was Q-TOF with an MS^2^ mass accuracy of 10ppm. All other settings were default for batch computation. CSI:FingerID predictions were calculated with Fallback adducts including [M-H]-, [M+H]+, [M+NH4]+, [M+Na]+, [M+Cl]-, [M+K]+, and [M-H2O+H]+ and search DBs was set to the Bio Database. Spectra for which CSI:FingerID returned no predicted structures were counted as not passing the requisite Tanimoto similarity threshold for the purposes of calculating accuracy.

### Isolation Experiments

#### General Procedures

All solvents and chemicals were purchased and used without further purification.

#### Complex mixture extract preparation

*Withania somnifera* primary extract (4g. standards.ONE) was fractionated by solid phase extraction through 20g HP20 resin (250 µm particle size) on a CombiFlash EZ-Prep (Teledyne) into 5 fractions (100 % H_2_O at 15 mL/min, 20% MeOH in H_2_O at 15 mL/min, 50% MeOH in H_2_O 35 mL/min, 80% MeOH in H_2_O at 35 ml/min, dichloromethane at 35 mL/min, 20 min for each fraction). The 50% MeOH fraction was evaporated and dried under vacuum to yield 323 mg of a dark oil. A 100 mg/mL solution of the dried fraction was prepared by redissolving with 50% MeOH in H_2_O. A mixture of the 26 endogenous natural products was prepared in MeOH through serial dilution to produce a solution containing ∼50 ng/100 µL of each compound. A 100 µL aliquot of this solution was then combined with a 100 µL aliquot of the 50% MeOH in H_2_O solution.

### LC-MS/MS analysis of the complex mixture

The 50% MeOH in H2O fraction spiked with the 26 exogenous natural products was profiled on a semi-prep LC system on a C18 column (5µm particle size, 10 x 250 mm, Waters Corporation) with a reversed-phase gradient (solvent A: CH_3_CN, solvent B: H2O, 2% A to 82% A over 45 min, 6 mL/minute) using a 200 µL sample injection. The solvent flow is then split 1:100 with the low output flow directed towards untargeted MS/MS analysis on Bruker timsTOF Pro 2 mass spectrometers (Bruker Daltonics, Billerica, MA). The low flow is made up with either 0.1% formic acid or 5mM ammonium formate in 90:10 water:acetonitrile mixture to enhance ionization efficiency for positive and negative heated electrospray ionization (HESI) respectively. The MS method used is designed for precise small molecule metabolomics, utilizing DDA-PASEF (data dependent acquisition - parallel accumulation serial fragmentation) TOF MS/MS with a 2 second precursor exclusion window, running at 10 Hz from 20 to 1300 Dalton, with CID energies ramping from 30-40 eV. The method uses HESI with a capillary voltage of 4500V, a dry gas temperature of 220 Celsius and sheath gas at 100 Celsius.

### General method for Isolation of endogenous compounds

Isolation of the nine endogenous compounds was conducted by reverse phase LCMS system (Agilent 1260 LC and MSD XT) with a 4.6mm x 250mm Luna Omega Polar C18 column (Phenomenex) using acetonitrile/water solvent gradient. The isolates were then identified by 1D and 2D-NMR data and comparison to published data. All NMR spectra (except Isolate C and H) were acquired on a Bruker Avance III 600 spectrometer and a Bruker Cryoplatform with 1.7mm inverse detection triple resonance cryoprobe with z-gradients. Isolate C and H were acquired on a Bruker Avance III 300 spectrometer with 5mm BBFO probe.

## ASSOCIATED CONTENT

### Supporting Information

Additional data on the extract set of compounds including structures, predicted properties, and calculated properties based on confirmed structure are found in the Supplemental Information, Table S1. This material is available free of charge at http://pubs.acs.org

## AUTHOR INFORMATION

## ACKNOWLEDGEMENTS

The authors would like to thank Joseph Pategou for assistance with Figs. 1, 7, and 8, Amy Myers for data research, and to A. Allen, D. Domingo-Fernandez, and K. Ajayi for manuscript review.

## Notes

### Competing Interest Statement

PCD is a Scientific Advisor to Cybele and a Scientific Co-founder and advisor to Ometa and Enveda with prior approval by UC-San Diego. All other authors are employees of Enveda.

### Summary of Updates

Manuscript substantially rewritten; New section added; Additional authors.

