## Supplemental Table 1 for "MS2Prop: A machine learning model that directly generates *de novo* predictions of drug-likeness of natural products from unannotated MS/MS spectra"

[‖]unaffilated

[⊥]Uncountable Innovations Consulting, Boise, ID, USA

Table Contents

Table S1: Extract dataset compounds, MS2Prop predictions, and calculated values

Extract dataset chemical structures

Isolate analytical data

Table S1: Extract dataset compounds, MS2Prop predictions, and calculated values

| Entry | 1 | 2 | | 3 | |
|---|---|---|---|---|---|
| name | (-)-Huperzine A | (20S)-Protopanaxatriol | | Carnosic acid | |
| Retention Time (min) | | | | | |
| annotated_adduct | **[M+H]+** | **[M+H-2H2O]+** | **[M+H-H2O]+** | **[M+H-2H2O]+** | **[M+H-H2O]+** |
| m/z | | | | | |
| m/z intensity | 742,054 | 85,782 | 50,871 | 199,749 | 248,658 |
| predicted alogp | 2.60 | 7.30 | 6.79 | 2.68 | 2.11 |
| actual alogp | 2.00 | 5.47 | 5.47 | 4.32 | 4.32 |
| error alogp | 0.60 | 1.83 | 1.32 | -1.64 | -2.21 |
| predicted HBA | 2.35 | 3.03 | 3.78 | 4.65 | 4.63 |
| actual HBA | 2 | 4 | 4 | 4 | 4 |
| error HBA | 0.35 | -0.97 | -0.22 | 0.65 | 0.63 |
| predicted HBD | 1.13 | 1.79 | 1.63 | 2.61 | 2.04 |
| actual HBD | 2 | 4 | 4 | 3 | 3 |
| error HBD | -0.87 | -2.21 | -2.37 | -0.39 | -0.96 |
| predicted TPSA | 41.85 | 53.33 | 68.82 | 91.55 | 76.33 |
| actual TPSA | 58.88 | 80.92 | 80.92 | 77.76 | 77.76 |
| error TPSA | -17.03 | -27.59 | -12.10 | 13.79 | -1.43 |
| predicted # rot. bonds | 1.45 | 10.45 | 10.29 | 9.57 | 6.11 |
| actual # rot. bonds | 0 | 4 | 4 | 2 | 2 |
| error # rot. Bonds | 1.45 | 6.45 | 6.29 | 7.57 | 4.11 |
| predicted # arom. rings | 0.72 | 0.11 | 0.18 | 0.00 | 0.34 |
| actual # arom. Rings | 1 | 0 | 0 | 1 | 1 |
| error # arom. Rings | -0.28 | 0.11 | 0.18 | -1.00 | -0.66 |
| predicted # aliph. rings | 1.97 | 2.28 | 3.04 | 0.63 | 1.87 |
| actual # aliph. Rings | 2 | 4 | 4 | 2 | 2 |
| error # aliph. Rings | -0.03 | -1.72 | -0.96 | -1.37 | -0.13 |
| predicted Fsp3 | 0.57 | 0.80 | 0.87 | 0.55 | 0.57 |
| actual Fsp3 | 0.40 | 0.93 | 0.93 | 0.65 | 0.65 |
| error Fsp3 | 0.17 | -0.13 | -0.06 | -0.10 | -0.08 |
| predicted QED | 0.77 | 0.32 | 0.30 | 0.36 | 0.58 |
| actual QED | 0.68 | 0.41 | 0.41 | 0.71 | 0.71 |
| error QED | 0.09 | -0.08 | -0.11 | -0.35 | -0.12 |
| predicted SA | 3.69 | 4.13 | 4.55 | 3.98 | 3.85 |
| actual SA | 4.94 | 5.05 | 5.05 | 3.77 | 3.77 |
| error SA | -1.25 | -0.93 | -0.50 | 0.21 | 0.08 |

Table S1(cont.): Extract dataset compounds, MS2Prop predictions, and calculated values

| Entry | 4 | 5 | 6 | 7 | |
|---|---|---|---|---|---|
| name | **Cephalomannine** | **Colchicine** | **Cycloastragenol** | Fraxinellone | |
| Retention Time (min) | | | | | |
| annotated_adduct | **[M+H]+** | **[M+H]+** | **[M+H-H2O]+** | **[M+H-H2O]+** | **[M+H]+** |
| m/z | | | | | |
| m/z intensity | 73,731 | 450,965 | 736,141 | 632,074 | 521,994 |
| predicted alogp | 4.72 | 2.33 | 6.74 | 2.51 | 2.24 |
| actual alogp | 3.39 | 2.87 | 4.44 | 3.38 | 3.38 |
| error alogp | 1.33 | -0.54 | 2.31 | -0.87 | -1.15 |
| predicted HBA | 14.01 | 6.08 | 3.92 | 2.71 | 3.02 |
| actual HBA | 14 | 6 | 5 | 3 | 3 |
| error HBA | 0.01 | 0.08 | -1.08 | -0.29 | 0.02 |
| predicted HBD | 2.36 | 2.42 | 1.93 | 0.66 | 0.57 |
| actual HBD | 4 | 1 | 4 | 0 | 0 |
| error HBD | -1.64 | 1.42 | -2.07 | 0.66 | 0.57 |
| predicted TPSA | 202.23 | 98.00 | 74.57 | 44.40 | 52.57 |
| actual TPSA | 221.29 | 83.09 | 90.15 | 39.44 | 39.44 |
| error TPSA | -19.06 | 14.91 | -15.58 | 4.96 | 13.13 |
| predicted # rot. bonds | 8.72 | 3.92 | 16.23 | 2.44 | 2.91 |
| actual # rot. bonds | 10 | 5 | 2 | 1 | 1 |
| error # rot. Bonds | -1.28 | -1.08 | 14.23 | 1.44 | 1.91 |
| predicted # arom. rings | 1.31 | 0.97 | 1.01 | 0.43 | 0.93 |
| actual # arom. Rings | 2 | 2 | 0 | 1 | 1 |
| error # arom. Rings | -0.69 | -1.03 | 1.01 | -0.57 | -0.07 |
| predicted # aliph. rings | 3.43 | 2.15 | 0.37 | 1.19 | 0.62 |
| actual # aliph. Rings | 4 | 1 | 6 | 2 | 2 |
| error # aliph. Rings | -0.57 | 1.15 | -5.63 | -0.81 | -1.38 |
| predicted Fsp3 | 0.60 | 0.51 | 0.68 | 0.44 | 0.30 |
| actual Fsp3 | 0.51 | 0.36 | 1.00 | 0.50 | 0.50 |
| error Fsp3 | 0.09 | 0.15 | -0.32 | -0.06 | -0.20 |
| predicted QED | 0.11 | 0.61 | 0.12 | 0.70 | 0.52 |
| actual QED | 0.12 | 0.83 | 0.46 | 0.70 | 0.70 |
| error QED | 0.00 | -0.22 | -0.34 | 0.01 | -0.18 |
| predicted SA | 6.50 | 4.53 | 3.35 | 3.01 | 2.77 |
| actual SA | 6.09 | 2.92 | 5.97 | 3.89 | 3.89 |
| error SA | 0.41 | 1.60 | -2.62 | -0.88 | -1.12 |

Table S1: Extract dataset compounds, MS2Prop predictions, and calculated values

| Entry | 8 | 9 | 10 | 11 | 12 |
|---|---|---|---|---|---|
| name | Galanthamine HBr | Ginkgolide A | Ginkgolide J | Huperzine B | Lycorine |
| Retention Time (min) | | | | | |
| annotated_adduct | **[M+H]+** | **[M+NH4]+** | **[M+NH4]+** | **[M+H]+** | **[M+H]+** |
| m/z | | | | | |
| m/z intensity | 693,131 | 128,840 | 32,139 | 595,125 | 760,514 |
| predicted alogp | 1.56 | 1.47 | 0.04 | 1.38 | 0.25 |
| actual alogp | 1.85 | -0.34 | -1.37 | 2.09 | 0.75 |
| error alogp | -0.29 | 1.81 | 1.41 | -0.72 | -0.50 |
| predicted HBA | 4.47 | 7.51 | 8.46 | 3.62 | 4.92 |
| actual HBA | 4 | 9 | 10 | 2 | 5 |
| error HBA | 0.47 | -1.49 | -1.54 | 1.62 | -0.08 |
| predicted HBD | 1.94 | 2.52 | 4.86 | 1.91 | 3.11 |
| actual HBD | 1 | 2 | 3 | 2 | 2 |
| error HBD | 0.94 | 0.52 | 1.86 | -0.09 | 1.11 |
| predicted TPSA | 77.13 | 117.00 | 148.24 | 59.43 | 91.27 |
| actual TPSA | 41.93 | 128.59 | 148.82 | 44.89 | 62.16 |
| error TPSA | 35.20 | -11.59 | -0.58 | 14.54 | 29.11 |
| predicted # rot. bonds | 6.65 | 4.92 | 5.64 | 3.81 | 4.05 |
| actual # rot. bonds | 1 | 0 | 0 | 0 | 0 |
| error # rot. Bonds | 5.65 | 4.92 | 5.64 | 3.81 | 4.05 |
| predicted # arom. rings | 0.79 | 0.79 | 1.48 | 0.81 | 0.92 |
| actual # arom. Rings | 1 | 0 | 0 | 1 | 1 |
| error # arom. Rings | -0.21 | 0.79 | 1.48 | -0.19 | -0.08 |
| predicted # aliph. rings | 0.54 | 2.86 | 2.13 | 1.45 | 1.40 |
| actual # aliph. Rings | 3 | 6 | 6 | 3 | 4 |
| error # aliph. Rings | -2.46 | -3.14 | -3.87 | -1.55 | -2.60 |
| predicted Fsp3 | 0.54 | 0.55 | 0.55 | 0.57 | 0.43 |
| actual Fsp3 | 0.53 | 0.85 | 0.85 | 0.56 | 0.50 |
| error Fsp3 | 0.01 | -0.30 | -0.30 | 0.01 | -0.07 |
| predicted QED | 0.61 | 0.52 | 0.36 | 0.59 | 0.61 |
| actual QED | 0.80 | 0.41 | 0.31 | 0.70 | 0.69 |
| error QED | -0.20 | 0.11 | 0.05 | -0.10 | -0.08 |
| predicted SA | 2.97 | 5.05 | 4.56 | 3.31 | 3.31 |
| actual SA | 4.23 | 6.24 | 6.28 | 5.00 | 4.17 |
| error SA | -1.26 | -1.18 | -1.72 | -1.70 | -0.85 |

Table S1(cont.): Extract dataset compounds, MS2Prop predictions, and calculated values

| Entry | 13 | 14 | 15 | 16 | |
|---|---|---|---|---|---|
| name | Gypsogenin-3-O-beta-D-glucuronide methyl ester | Monocrotaline | Nardosinone | Panaxtriol | |
| Retention Time (min) | | | | | |
| annotated_adduct | **[M+NH4]+** | **[M+H]+** | **[M+H-H2O]+** | **[M+H-H2O]+** | **[M+H]+** |
| m/z | | | | | |
| m/z intensity | 24,261 | 584,717 | 464,855 | 2,618 | 800,729 |
| predicted alogp | 1.42 | 0.61 | 2.20 | 7.00 | 5.26 |
| actual alogp | 4.42 | -0.39 | 3.05 | 5.71 | 5.71 |
| error alogp | -2.99 | 1.00 | -0.85 | 1.29 | -0.45 |
| predicted HBA | 14.02 | 5.89 | 3.11 | 2.67 | 3.81 |
| actual HBA | 10 | 7 | 3 | 4 | 4 |
| error HBA | 4.02 | -1.11 | 0.11 | -1.33 | -0.19 |
| predicted HBD | 6.96 | 2.43 | 2.13 | 1.54 | 2.93 |
| actual HBD | 4 | 2 | 0 | 3 | 3 |
| error HBD | 2.96 | 0.43 | 2.13 | -1.46 | -0.07 |
| predicted TPSA | 238.06 | 101.86 | 62.50 | 54.54 | 70.31 |
| actual TPSA | 159.82 | 96.30 | 35.53 | 69.92 | 69.92 |
| error TPSA | 78.24 | 5.56 | 26.97 | -15.38 | 0.39 |
| predicted # rot. bonds | 20.58 | 2.36 | 5.05 | 2.72 | 2.90 |
| actual # rot. bonds | 5 | 0 | 0 | 1 | 1 |
| error # rot. Bonds | 15.58 | 2.36 | 5.05 | 1.72 | 1.90 |
| predicted # arom. rings | 0.07 | 0.06 | 0.24 | -0.06 | 0.15 |
| actual # arom. Rings | 0 | 0 | 0 | 0 | 0 |
| error # arom. Rings | 0.07 | 0.06 | 0.24 | -0.06 | 0.15 |
| predicted # aliph. rings | 0.41 | 2.55 | 1.27 | 4.87 | 4.60 |
| actual # aliph. Rings | 6 | 3 | 3 | 5 | 5 |
| error # aliph. Rings | -5.59 | -0.45 | -1.73 | -0.13 | -0.40 |
| predicted Fsp3 | 0.69 | 0.72 | 0.62 | 0.91 | 0.94 |
| actual Fsp3 | 0.86 | 0.75 | 0.80 | 1.00 | 1.00 |
| error Fsp3 | -0.18 | -0.03 | -0.18 | -0.09 | -0.06 |
| predicted QED | 0.06 | 0.45 | 0.54 | 0.38 | 0.47 |
| actual QED | 0.14 | 0.46 | 0.62 | 0.45 | 0.45 |
| error QED | -0.09 | -0.01 | -0.08 | -0.07 | 0.01 |
| predicted SA | 5.08 | 4.92 | 3.83 | 4.77 | 5.23 |
| actual SA | 5.44 | 5.04 | 4.80 | 5.06 | 5.06 |
| error SA | -0.36 | -0.12 | -0.97 | -0.29 | 0.17 |

Table S1: Extract dataset compounds, MS2Prop predictions, and calculated values

| Entry | 17 | | | 18 | |
|---|---|---|---|---|---|
| name | **Parthenolide** | | | **Phorbol** | |
| Retention Time (min) | | | | | |
| annotated_adduct | **[M+H-2H2O]+** | **[M+H-H2O]+** | **[M+H]+** | **[M+H-2H2O]+** | **[M+H-3H2O]+** |
| m/z | | | | | |
| m/z intensity | 142,524 | 1,048,911 | 183,017 | 168,215 | 148,379 |
| predicted alogp | 2.49 | 3.01 | 1.62 | 0.57 | 1.42 |
| actual alogp | 2.76 | 2.76 | 2.76 | -0.07 | -0.07 |
| error alogp | -0.27 | 0.25 | -1.14 | 0.64 | 1.49 |
| predicted HBA | 2.18 | 2.25 | 3.81 | 5.23 | 4.60 |
| actual HBA | 3 | 3 | 3 | 6 | 6 |
| error HBA | -0.82 | -0.75 | 0.81 | -0.77 | -1.40 |
| predicted HBD | 0.96 | 0.37 | 1.81 | 3.36 | 2.62 |
| actual HBD | 0 | 0 | 0 | 5 | 5 |
| error HBD | 0.96 | 0.37 | 1.81 | -1.64 | -2.38 |
| predicted TPSA | 44.19 | 38.05 | 71.96 | 98.23 | 80.01 |
| actual TPSA | 38.83 | 38.83 | 38.83 | 118.22 | 118.22 |
| error TPSA | 5.36 | -0.78 | 33.13 | -19.99 | -38.21 |
| predicted # rot. bonds | 1.10 | 0.69 | 2.63 | 2.55 | 2.99 |
| actual # rot. bonds | 0 | 0 | 0 | 1 | 1 |
| error # rot. Bonds | 1.10 | 0.69 | 2.63 | 1.55 | 1.99 |
| predicted # arom. rings | 2.00 | 0.93 | 0.48 | 0.19 | 0.66 |
| actual # arom. Rings | 0 | 0 | 0 | 0 | 0 |
| error # arom. Rings | 2.00 | 0.93 | 0.48 | 0.19 | 0.66 |
| predicted # aliph. rings | 0.21 | 1.61 | 1.57 | 3.99 | 3.49 |
| actual # aliph. Rings | 3 | 3 | 3 | 4 | 4 |
| error # aliph. Rings | -2.79 | -1.39 | -1.43 | -0.01 | -0.51 |
| predicted Fsp3 | 0.18 | 0.43 | 0.49 | 0.70 | 0.61 |
| actual Fsp3 | 0.67 | 0.67 | 0.67 | 0.75 | 0.75 |
| error Fsp3 | -0.49 | -0.24 | -0.17 | -0.05 | -0.14 |
| predicted QED | 0.58 | 0.52 | 0.64 | 0.55 | 0.64 |
| actual QED | 0.29 | 0.29 | 0.29 | 0.42 | 0.42 |
| error QED | 0.29 | 0.24 | 0.35 | 0.14 | 0.23 |
| predicted SA | 2.05 | 3.33 | 3.44 | 5.75 | 5.76 |
| actual SA | 4.72 | 4.72 | 4.72 | 5.18 | 5.18 |
| error SA | -2.67 | -1.39 | -1.28 | 0.57 | 0.58 |

Table S1(cont.): Extract dataset compounds, MS2Prop predictions, and calculated values

| Entry | 19 | | 20 | 21 | |
|---|---|---|---|---|---|
| name | **Piperine** | | **Piperlongumine** | **Piperlonguminine** | |
| Retention Time (min) | | | | | |
| annotated_adduct | **[2M+H]+** | **[M+H]+** | **[M+H]+** | **[2M+H]+** | **[M+H]+** |
| m/z | | | | | |
| m/z intensity | 51,862 | 1,340,739 | 1,518,055 | 222,765 | 2,271,027 |
| predicted alogp | 2.60 | 2.40 | 2.12 | 6.95 | 3.09 |
| actual alogp | 3.00 | 3.00 | 2.04 | 2.76 | 2.76 |
| error alogp | -0.40 | -0.60 | 0.08 | 4.19 | 0.33 |
| predicted HBA | 8.40 | 3.60 | 5.09 | 5.99 | 3.35 |
| actual HBA | 3 | 3 | 5 | 3 | 3 |
| error HBA | 5.40 | 0.60 | 0.09 | 2.99 | 0.35 |
| predicted HBD | 2.85 | 0.20 | 0.22 | 2.51 | 1.04 |
| actual HBD | 0 | 0 | 0 | 1 | 1 |
| error HBD | 2.85 | 0.20 | 0.22 | 1.51 | 0.04 |
| predicted TPSA | 140.63 | 49.46 | 70.71 | 108.61 | 53.51 |
| actual TPSA | 38.77 | 38.77 | 65.07 | 47.56 | 47.56 |
| error TPSA | 101.86 | 10.69 | 5.64 | 61.05 | 5.95 |
| predicted # rot. bonds | 5.17 | 3.09 | 5.14 | 14.29 | 3.25 |
| actual # rot. bonds | 3 | 3 | 5 | 5 | 5 |
| error # rot. Bonds | 2.17 | 0.09 | 0.14 | 9.29 | -1.75 |
| predicted # arom. rings | 0.02 | 1.22 | 1.00 | 2.98 | 1.87 |
| actual # arom. Rings | 1 | 1 | 1 | 1 | 1 |
| error # arom. Rings | -0.98 | 0.22 | 0.00 | 1.98 | 0.87 |
| predicted # aliph. rings | 3.71 | 1.42 | 0.79 | 0.57 | 0.41 |
| actual # aliph. Rings | 2 | 2 | 1 | 1 | 1 |
| error # aliph. Rings | 1.71 | -0.58 | -0.21 | -0.43 | -0.59 |
| predicted Fsp3 | 0.70 | 0.34 | 0.37 | 0.40 | 0.24 |
| actual Fsp3 | 0.35 | 0.35 | 0.29 | 0.31 | 0.31 |
| error Fsp3 | 0.35 | -0.01 | 0.08 | 0.09 | -0.07 |
| predicted QED | 0.28 | 0.69 | 0.75 | 0.12 | 0.65 |
| actual QED | 0.63 | 0.63 | 0.78 | 0.66 | 0.66 |
| error QED | -0.35 | 0.06 | -0.03 | -0.54 | -0.01 |
| predicted SA | 5.95 | 2.26 | 2.42 | 3.34 | 2.44 |
| actual SA | 2.34 | 2.34 | 2.59 | 2.40 | 2.40 |
| error SA | 3.60 | -0.08 | -0.17 | 0.94 | 0.04 |

Table S1: Extract dataset compounds, MS2Prop predictions, and calculated values

| Entry | 22 | 23 | 24 | 25 | 26 |
|---|---|---|---|---|---|
| name | **Pseudolaric Acid B** | **Sophocarpine** | **Sophoridine** | **Tenuifolin** | **Tetrandrine** |
| Retention Time (min) | | | | | |
| annotated_adduct | **[M+NH4]+** | **[M+H]+** | **[M+H]+** | **[M+NH4]+** | **[M+Na]+** |
| m/z | | | | | |
| m/z intensity | 189,650 | 1,118,375 | 432,721 | 190,667 | 394,091 |
| predicted alogp | 3.68 | 2.01 | 2.00 | 3.27 | 2.50 |
| actual alogp | 2.87 | 1.65 | 1.87 | 2.07 | 7.16 |
| error alogp | 0.81 | 0.36 | 0.13 | 1.21 | -4.67 |
| predicted HBA | 7.01 | 2.56 | 3.54 | 10.64 | 10.45 |
| actual HBA | 8 | 2 | 2 | 12 | 8 |
| error HBA | -0.99 | 0.56 | 1.54 | -1.36 | 2.45 |
| predicted HBD | 1.05 | 0.67 | 1.70 | 5.92 | 5.17 |
| actual HBD | 1 | 0 | 0 | 8 | 0 |
| error HBD | 0.05 | 0.67 | 1.70 | -2.08 | 5.17 |
| predicted TPSA | 97.58 | 38.15 | 63.13 | 184.98 | 194.75 |
| actual TPSA | 116.20 | 23.55 | 23.55 | 214.44 | 61.86 |
| error TPSA | -18.62 | 14.60 | 39.58 | -29.46 | 132.89 |
| predicted # rot. bonds | 6.12 | 1.74 | 7.58 | 9.26 | 10.47 |
| actual # rot. bonds | 5 | 0 | 0 | 6 | 4 |
| error # rot. Bonds | 1.12 | 1.74 | 7.58 | 3.26 | 6.47 |
| predicted # arom. rings | 1.14 | 0.18 | 0.14 | 0.39 | 0.23 |
| actual # arom. Rings | 0 | 0 | 0 | 0 | 4 |
| error # arom. Rings | 1.14 | 0.18 | 0.14 | 0.39 | -3.77 |
| predicted # aliph. rings | 1.55 | 2.87 | 0.50 | 4.57 | 3.55 |
| actual # aliph. Rings | 3 | 4 | 4 | 6 | 4 |
| error # aliph. Rings | -1.45 | -1.13 | -3.50 | -1.43 | -0.45 |
| predicted Fsp3 | 0.53 | 0.73 | 0.66 | 0.76 | 0.66 |
| actual Fsp3 | 0.57 | 0.80 | 0.93 | 0.89 | 0.37 |
| error Fsp3 | -0.04 | -0.07 | -0.27 | -0.13 | 0.29 |
| predicted QED | 0.53 | 0.71 | 0.47 | 0.11 | 0.12 |
| actual QED | 0.31 | 0.65 | 0.65 | 0.15 | 0.24 |
| error QED | 0.23 | 0.06 | -0.19 | -0.04 | -0.11 |
| predicted SA | 4.40 | 4.11 | 3.38 | 5.39 | 5.84 |
| actual SA | 5.33 | 4.11 | 3.71 | 5.59 | 5.73 |
| error SA | -0.93 | 0.00 | -0.33 | -0.21 | 0.12 |

Table S1(cont.): Extract dataset compounds, MS2Prop predictions, and calculated values

| Entry | 27 | | | 28 |
|---|---|---|---|---|
| name | Isolate A | | | Isolate B |
| Retention Time (min) | | | | |
| annotated_adduct | **[2M+NH4]+** | **[M+H]+** | **[M+NH4]+** | **[M+H]+** |
| m/z | | | | |
| m/z intensity | 187,626 | 4,092,922 | 14,307,520 | 4,461,692 |
| predicted alogp | 4.29 | 4.33 | 4.59 | 0.90 |
| actual alogp | 3.35 | 3.35 | 3.35 | 0.97 |
| error alogp | 0.94 | 0.97 | 1.24 | -0.07 |
| predicted HBA | 12.69 | 5.37 | 5.95 | 5.11 |
| actual HBA | 6 | 6 | 6 | 4 |
| error HBA | 6.69 | -0.63 | -0.05 | 1.11 |
| predicted HBD | 4.26 | 1.50 | 2.25 | 3.44 |
| actual HBD | 2 | 2 | 2 | 4 |
| error HBD | 2.26 | -0.50 | 0.25 | -0.56 |
| predicted TPSA | 219.06 | 87.30 | 107.13 | 107.41 |
| actual TPSA | 96.36 | 96.36 | 96.36 | 95.58 |
| error TPSA | 122.70 | -9.06 | 10.77 | 11.83 |
| predicted # rot. bonds | 7.58 | 3.91 | 7.99 | 4.47 |
| actual # rot. bonds | 3 | 3 | 3 | 6 |
| error # rot. Bonds | 4.58 | 0.91 | 4.99 | -1.53 |
| predicted # arom. rings | 0.31 | 0.10 | 0.16 | 1.08 |
| actual # arom. Rings | 0 | 0 | 0 | 1 |
| error # arom. Rings | 0.31 | 0.10 | 0.16 | 0.08 |
| predicted # aliph. rings | 2.19 | 4.94 | 2.95 | -0.14 |
| actual # aliph. Rings | 6 | 6 | 6 | 0 |
| error # aliph. Rings | -3.81 | -1.06 | -3.05 | -0.14 |
| predicted Fsp3 | 0.69 | 0.76 | 0.62 | 0.16 |
| actual Fsp3 | 0.79 | 0.79 | 0.79 | 0.31 |
| error Fsp3 | -0.10 | -0.03 | -0.16 | -0.15 |
| predicted QED | 0.17 | 0.38 | 0.28 | 0.40 |
| actual QED | 0.49 | 0.49 | 0.49 | 0.34 |
| error QED | -0.32 | -0.11 | -0.21 | 0.05 |
| predicted SA | 6.71 | 5.38 | 5.46 | 2.32 |
| actual SA | 5.72 | 5.72 | 5.72 | 2.17 |
| error SA | 0.99 | -0.34 | -0.26 | 0.16 |

Table S1: Extract dataset compounds, MS2Prop predictions, and calculated values

| Entry | 29 | | | | 30 |
|---|---|---|---|---|---|
| name | Isolate C | | | | Isolate D |
| Retention Time (min) | | | | | |
| annotated_adduct | **[M+H]+** | **[M+K]+** | **[M+NH4]+** | **[M-H+2K]+** | **[M+H]+** |
| m/z | | | | | |
| m/z intensity | 726,513 | 311,176 | 9,865,723 | 225,204 | 292,012 |
| predicted alogp | 3.21 | 4.31 | 4.73 | 2.18 | 2.37 |
| actual alogp | 2.37 | 2.37 | 2.37 | 2.37 | 1.27 |
| error alogp | 0.84 | 1.94 | 2.35 | -0.19 | 1.10 |
| predicted HBA | 9.78 | 9.64 | 8.91 | 11.70 | 2.54 |
| actual HBA | 10 | 10 | 10 | 10 | 3 |
| error HBA | -0.22 | -0.36 | -1.09 | 1.70 | -0.46 |
| predicted HBD | 2.31 | 1.86 | 1.87 | 2.02 | 0.41 |
| actual HBD | 3 | 3 | 3 | 3 | 3 |
| error HBD | -0.69 | -1.14 | -1.13 | -0.98 | -2.59 |
| predicted TPSA | 147.05 | 142.32 | 131.57 | 169.16 | 41.66 |
| actual TPSA | 159.96 | 159.96 | 159.96 | 159.96 | 65.12 |
| error TPSA | -12.91 | -17.64 | -28.39 | 9.20 | -23.46 |
| predicted # rot. bonds | 6.14 | 6.56 | 7.46 | 4.61 | 1.06 |
| actual # rot. bonds | 5 | 5 | 5 | 5 | 1 |
| error # rot. Bonds | 1.14 | 1.56 | 2.46 | -0.39 | 0.06 |
| predicted # arom. rings | 0.61 | 0.42 | 0.33 | 0.17 | 0.22 |
| actual # arom. Rings | 0 | 0 | 0 | 0 | 2 |
| error # arom. Rings | 0.61 | 0.42 | 0.33 | 0.17 | -1.78 |
| predicted # aliph. rings | 4.07 | 4.43 | 4.18 | 5.15 | 1.60 |
| actual # aliph. Rings | 6 | 6 | 6 | 6 | 1 |
| error # aliph. Rings | -1.93 | -1.57 | -1.82 | -0.85 | 0.60 |
| predicted Fsp3 | 0.59 | 0.67 | 0.71 | 0.65 | 0.58 |
| actual Fsp3 | 0.86 | 0.86 | 0.86 | 0.86 | 0.25 |
| error Fsp3 | -0.27 | -0.19 | -0.14 | -0.21 | 0.33 |
| predicted QED | 0.19 | 0.21 | 0.20 | 0.17 | 0.58 |
| actual QED | 0.26 | 0.26 | 0.26 | 0.26 | 0.67 |
| error QED | -0.06 | -0.05 | -0.06 | -0.08 | -0.10 |
| predicted SA | 5.94 | 5.79 | 5.73 | 6.24 | 3.47 |
| actual SA | 5.94 | 5.94 | 5.94 | 5.94 | 2.88 |
| error SA | 0.01 | -0.15 | -0.21 | 0.31 | 0.59 |

Table S1(cont.): Extract dataset compounds, MS2Prop predictions, and calculated values

| Entry | 31 | | | | |
|---|---|---|---|---|---|
| name | Isolate E | | | | |
| Retention Time (min) | | | | | |
| annotated_adduct | **[M+H+DMSO]+** | **[M+H-3H2O]+** | **[M+H-H2O]+** | **[M+H]+** | **[M+NH4]+** |
| m/z | | | | | |
| m/z intensity | 494,821 | 15,742 | 33,185 | 88,072 | 4,438,231 |
| predicted alogp | 1.71 | 3.10 | 3.99 | 1.00 | 2.92 |
| actual alogp | 3.50 | 3.50 | 3.50 | 3.50 | 3.50 |
| error alogp | -1.79 | -0.40 | 0.50 | -2.50 | -0.57 |
| predicted HBA | 9.00 | 6.82 | 5.79 | 7.68 | 6.73 |
| actual HBA | 6 | 6 | 6 | 6 | 6 |
| error HBA | 3.00 | 0.82 | -0.21 | 1.68 | 0.73 |
| predicted HBD | 4.92 | 2.98 | 3.06 | 6.22 | 3.43 |
| actual HBD | 2 | 2 | 2 | 2 | 2 |
| error HBD | 2.92 | 0.98 | 1.06 | 4.22 | 1.43 |
| predicted TPSA | 165.36 | 112.22 | 109.01 | 164.22 | 126.35 |
| actual TPSA | 96.36 | 96.36 | 96.36 | 96.36 | 96.36 |
| error TPSA | 69.00 | 15.86 | 12.65 | 67.86 | 29.99 |
| predicted # rot. bonds | 9.61 | 16.14 | 12.00 | 6.74 | 7.38 |
| actual # rot. bonds | 2 | 2 | 2 | 2 | 2 |
| error # rot. Bonds | 7.61 | 14.14 | 10.00 | 4.74 | 5.38 |
| predicted # arom. rings | 0.00 | 0.32 | 0.18 | 0.07 | 0.34 |
| actual # arom. Rings | 0 | 0 | 0 | 0 | 0 |
| error # arom. Rings | 0.00 | 0.32 | 0.18 | 0.07 | 0.34 |
| predicted # aliph. rings | 3.11 | 0.19 | 1.34 | 2.50 | 3.63 |
| actual # aliph. Rings | 6 | 6 | 6 | 6 | 6 |
| error # aliph. Rings | -2.89 | -5.81 | -4.66 | -3.50 | -2.37 |
| predicted Fsp3 | 0.75 | 0.76 | 0.62 | 0.76 | 0.69 |
| actual Fsp3 | 0.79 | 0.79 | 0.79 | 0.79 | 0.79 |
| error Fsp3 | -0.03 | -0.02 | -0.16 | -0.03 | -0.09 |
| predicted QED | 0.17 | 0.20 | 0.23 | 0.25 | 0.31 |
| actual QED | 0.47 | 0.47 | 0.47 | 0.47 | 0.47 |
| error QED | -0.30 | -0.27 | -0.24 | -0.23 | -0.17 |
| predicted SA | 5.09 | 3.47 | 4.69 | 5.29 | 5.91 |
| actual SA | 5.76 | 5.76 | 5.76 | 5.76 | 5.76 |
| error SA | -0.67 | -2.29 | -1.07 | -0.47 | 0.15 |

Table S1: Extract dataset compounds, MS2Prop predictions, and calculated values

| Entry | 32 | | 33 | | |
|---|---|---|---|---|---|
| name | **Isolate F** | | **Isolate G** | | |
| Retention Time (min) | | | | | |
| annotated_adduct | **[M+H]+** | **[M+NH4]+** | **[M+H-2H2O]+** | **[M+H-H2O]+** | **[M+NH4]+** |
| m/z | | | | | |
| m/z intensity | 7,351,391 | 8,303,941 | 42,684 | 22,174 | 2,668,464 |
| predicted alogp | 2.68 | 3.23 | 4.26 | 5.00 | 2.70 |
| actual alogp | 3.34 | 3.34 | 3.28 | 3.28 | 3.28 |
| error alogp | -0.67 | -0.12 | 0.98 | 1.72 | -0.58 |
| predicted HBA | 7.52 | 7.26 | 5.69 | 5.55 | 8.46 |
| actual HBA | 7 | 7 | 7 | 7 | 7 |
| error HBA | 0.52 | 0.26 | -1.31 | -1.45 | 1.46 |
| predicted HBD | 4.15 | 2.41 | 3.19 | 2.44 | 4.47 |
| actual HBD | 2 | 2 | 2 | 2 | 2 |
| error HBD | 2.15 | 0.41 | 1.19 | 0.44 | 2.47 |
| predicted TPSA | 131.46 | 119.87 | 107.22 | 91.69 | 146.56 |
| actual TPSA | 105.59 | 105.59 | 105.59 | 105.59 | 105.59 |
| error TPSA | 25.87 | 14.28 | 1.63 | -13.90 | 40.97 |
| predicted # rot. bonds | 4.56 | 5.68 | 5.22 | 1.95 | 9.29 |
| actual # rot. bonds | 3 | 3 | 3 | 3 | 3 |
| error # rot. Bonds | 1.56 | 2.68 | 2.22 | -1.05 | 6.29 |
| predicted # arom. rings | 0.04 | -0.13 | 0.00 | -0.01 | 0.46 |
| actual # arom. Rings | 0 | 0 | 0 | 0 | 0 |
| error # arom. Rings | 0.04 | -0.13 | 0.00 | -0.01 | 0.46 |
| predicted # aliph. rings | 4.67 | 3.88 | 3.65 | 5.27 | 3.00 |
| actual # aliph. Rings | 6 | 6 | 6 | 6 | 6 |
| error # aliph. Rings | -1.33 | -2.12 | -2.35 | -0.73 | -3.00 |
| predicted Fsp3 | 0.85 | 0.71 | 0.74 | 0.84 | 0.76 |
| actual Fsp3 | 0.86 | 0.86 | 0.93 | 0.93 | 0.93 |
| error Fsp3 | -0.02 | -0.15 | -0.20 | -0.09 | -0.17 |
| predicted QED | 0.34 | 0.27 | 0.36 | 0.38 | 0.26 |
| actual QED | 0.45 | 0.45 | 0.45 | 0.45 | 0.45 |
| error QED | -0.11 | -0.18 | -0.09 | -0.06 | -0.19 |
| predicted SA | 5.33 | 5.41 | 5.28 | 5.90 | 5.25 |
| actual SA | 5.88 | 5.88 | 6.04 | 6.04 | 6.04 |
| error SA | -0.55 | -0.47 | -0.76 | -0.14 | -0.79 |

Table S1(cont.): Extract dataset compounds, MS2Prop predictions, and calculated values

| Entry | 34 | | 35 |
|---|---|---|---|
| name | Isolate H | | Isolate I |
| Retention Time (min) | | | |
| annotated_adduct | **[M+H]+** | **[M+NH4]+** | **[M+H]+** |
| m/z | | | |
| m/z intensity | 20,546 | 289,153 | 1,153,156 |
| predicted alogp | 5.85 | 2.80 | 2.50 |
| actual alogp | 4.38 | 4.38 | 4.23 |
| error alogp | 1.47 | -1.58 | -1.73 |
| predicted HBA | 4.96 | 6.78 | 6.61 |
| actual HBA | 5 | 5 | 6 |
| error HBA | -0.04 | 1.78 | 0.61 |
| predicted HBD | 1.78 | 4.08 | 4.70 |
| actual HBD | 1 | 1 | 1 |
| error HBD | 0.78 | 3.08 | 3.70 |
| predicted TPSA | 80.50 | 126.95 | 122.20 |
| actual TPSA | 76.13 | 76.13 | 85.36 |
| error TPSA | 4.37 | 50.82 | 36.84 |
| predicted # rot. bonds | 8.41 | 8.71 | 3.53 |
| actual # rot. bonds | 2 | 2 | 3 |
| error # rot. Bonds | 6.41 | 6.71 | 0.53 |
| predicted # arom. rings | 0.02 | 0.16 | -0.03 |
| actual # arom. Rings | 0 | 0 | 0 |
| error # arom. Rings | 0.02 | 0.16 | -0.03 |
| predicted # aliph. rings | 1.44 | 3.06 | 5.01 |
| actual # aliph. Rings | 6 | 6 | 6 |
| error # aliph. Rings | -4.56 | -2.94 | -0.99 |
| predicted Fsp3 | 0.63 | 0.76 | 0.84 |
| actual Fsp3 | 0.79 | 0.79 | 0.86 |
| error Fsp3 | -0.16 | -0.02 | -0.03 |
| predicted QED | 0.32 | 0.29 | 0.35 |
| actual QED | 0.49 | 0.49 | 0.47 |
| error QED | -0.17 | -0.21 | -0.12 |
| predicted SA | 4.78 | 5.71 | 5.81 |
| actual SA | 5.63 | 5.63 | 5.76 |
| error SA | -0.86 | 0.07 | 0.05 |

Table S1: Extract dataset compounds, MS2Prop predictions, and calculated values

| Entry | 1 | 2 | | 3 | |
|---|---|---|---|---|---|
| name | (-)-Huperzine A | (20S)-Protopanaxatriol | | Carnosic acid | |
| Retention Time (min) | | | | | |
| annotated_adduct | **[M+H]+** | **[M+H-2H2O]+** | **[M+H-H2O]+** | **[M+H-2H2O]+** | **[M+H-H2O]+** |
| m/z | | | | | |
| m/z intensity | 742,054 | 85,782 | 50,871 | 199,749 | 248,658 |
| predicted alogp | 2.60 | 7.30 | 6.79 | 2.68 | 2.11 |
| actual alogp | 2.00 | 5.47 | 5.47 | 4.32 | 4.32 |
| error alogp | 0.60 | 1.83 | 1.32 | -1.64 | -2.21 |
| predicted HBA | 2.35 | 3.03 | 3.78 | 4.65 | 4.63 |
| actual HBA | 2 | 4 | 4 | 4 | 4 |
| error HBA | 0.35 | -0.97 | -0.22 | 0.65 | 0.63 |
| predicted HBD | 1.13 | 1.79 | 1.63 | 2.61 | 2.04 |
| actual HBD | 2 | 4 | 4 | 3 | 3 |
| error HBD | -0.87 | -2.21 | -2.37 | -0.39 | -0.96 |
| predicted TPSA | 41.85 | 53.33 | 68.82 | 91.55 | 76.33 |
| actual TPSA | 58.88 | 80.92 | 80.92 | 77.76 | 77.76 |
| error TPSA | -17.03 | -27.59 | -12.10 | 13.79 | -1.43 |
| predicted # rot. bonds | 1.45 | 10.45 | 10.29 | 9.57 | 6.11 |
| actual # rot. bonds | 0 | 4 | 4 | 2 | 2 |
| error # rot. Bonds | 1.45 | 6.45 | 6.29 | 7.57 | 4.11 |
| predicted # arom. rings | 0.72 | 0.11 | 0.18 | 0.00 | 0.34 |
| actual # arom. Rings | 1 | 0 | 0 | 1 | 1 |
| error # arom. Rings | -0.28 | 0.11 | 0.18 | -1.00 | -0.66 |
| predicted # aliph. rings | 1.97 | 2.28 | 3.04 | 0.63 | 1.87 |
| actual # aliph. Rings | 2 | 4 | 4 | 2 | 2 |
| error # aliph. Rings | -0.03 | -1.72 | -0.96 | -1.37 | -0.13 |
| predicted Fsp3 | 0.57 | 0.80 | 0.87 | 0.55 | 0.57 |
| actual Fsp3 | 0.40 | 0.93 | 0.93 | 0.65 | 0.65 |
| error Fsp3 | 0.17 | -0.13 | -0.06 | -0.10 | -0.08 |
| predicted QED | 0.77 | 0.32 | 0.30 | 0.36 | 0.58 |
| actual QED | 0.68 | 0.41 | 0.41 | 0.71 | 0.71 |
| error QED | 0.09 | -0.08 | -0.11 | -0.35 | -0.12 |
| predicted SA | 3.69 | 4.13 | 4.55 | 3.98 | 3.85 |
| actual SA | 4.94 | 5.05 | 5.05 | 3.77 | 3.77 |
| error SA | -1.25 | -0.93 | -0.50 | 0.21 | 0.08 |

Table S1(cont.): Extract dataset compounds, MS2Prop predictions, and calculated values

| |
|---|
| Entry |
| name |
| Retention Time (min) |
| annotated_adduct |
| m/z |
| m/z intensity |
| predicted alogp |
| actual alogp |
| error alogp |
| predicted HBA |
| actual HBA |
| error HBA |
| predicted HBD |
| actual HBD |
| error HBD |
| predicted TPSA |
| actual TPSA |
| error TPSA |
| predicted # rot. bonds |
| actual # rot. bonds |
| error # rot. Bonds |
| predicted # arom. rings |
| actual # arom. Rings |
| error # arom. Rings |
| predicted # aliph. rings |
| actual # aliph. Rings |
| error # aliph. Rings |
| predicted Fsp3 |
| actual Fsp3 |
| error Fsp3 |
| predicted QED |
| actual QED |
| error QED |
| predicted SA |
| actual SA |
| error SA |

Table S1: Extract dataset compounds, MS2Prop predictions, and calculated values

| Entry | 1 | 2 | | 3 | |
|---|---|---|---|---|---|
| name | (-)-Huperzine A | (20S)-Protopanaxatriol | | Carnosic acid | |
| Retention Time (min) | | | | | |
| annotated_adduct | **[M+H]+** | **[M+H-2H2O]+** | **[M+H-H2O]+** | **[M+H-2H2O]+** | **[M+H-H2O]+** |
| m/z | | | | | |
| m/z intensity | 742,054 | 85,782 | 50,871 | 199,749 | 248,658 |
| predicted alogp | 2.60 | 7.30 | 6.79 | 2.68 | 2.11 |
| actual alogp | 2.00 | 5.47 | 5.47 | 4.32 | 4.32 |
| error alogp | 0.60 | 1.83 | 1.32 | -1.64 | -2.21 |
| predicted HBA | 2.35 | 3.03 | 3.78 | 4.65 | 4.63 |
| actual HBA | 2 | 4 | 4 | 4 | 4 |
| error HBA | 0.35 | -0.97 | -0.22 | 0.65 | 0.63 |
| predicted HBD | 1.13 | 1.79 | 1.63 | 2.61 | 2.04 |
| actual HBD | 2 | 4 | 4 | 3 | 3 |
| error HBD | -0.87 | -2.21 | -2.37 | -0.39 | -0.96 |
| predicted TPSA | 41.85 | 53.33 | 68.82 | 91.55 | 76.33 |
| actual TPSA | 58.88 | 80.92 | 80.92 | 77.76 | 77.76 |
| error TPSA | -17.03 | -27.59 | -12.10 | 13.79 | -1.43 |
| predicted # rot. bonds | 1.45 | 10.45 | 10.29 | 9.57 | 6.11 |
| actual # rot. bonds | 0 | 4 | 4 | 2 | 2 |
| error # rot. Bonds | 1.45 | 6.45 | 6.29 | 7.57 | 4.11 |
| predicted # arom. rings | 0.72 | 0.11 | 0.18 | 0.00 | 0.34 |
| actual # arom. Rings | 1 | 0 | 0 | 1 | 1 |
| error # arom. Rings | -0.28 | 0.11 | 0.18 | -1.00 | -0.66 |
| predicted # aliph. rings | 1.97 | 2.28 | 3.04 | 0.63 | 1.87 |
| actual # aliph. Rings | 2 | 4 | 4 | 2 | 2 |
| error # aliph. Rings | -0.03 | -1.72 | -0.96 | -1.37 | -0.13 |
| predicted Fsp3 | 0.57 | 0.80 | 0.87 | 0.55 | 0.57 |
| actual Fsp3 | 0.40 | 0.93 | 0.93 | 0.65 | 0.65 |
| error Fsp3 | 0.17 | -0.13 | -0.06 | -0.10 | -0.08 |
| predicted QED | 0.77 | 0.32 | 0.30 | 0.36 | 0.58 |
| actual QED | 0.68 | 0.41 | 0.41 | 0.71 | 0.71 |
| error QED | 0.09 | -0.08 | -0.11 | -0.35 | -0.12 |
| predicted SA | 3.69 | 4.13 | 4.55 | 3.98 | 3.85 |
| actual SA | 4.94 | 5.05 | 5.05 | 3.77 | 3.77 |
| error SA | -1.25 | -0.93 | -0.50 | 0.21 | 0.08 |

Table S1(cont.): Extract dataset compounds, MS2Prop predictions, and calculated values

| |
|---|
| Entry |
| name |
| Retention Time (min) |
| annotated_adduct |
| m/z |
| m/z intensity |
| predicted alogp |
| actual alogp |
| error alogp |
| predicted HBA |
| actual HBA |
| error HBA |
| predicted HBD |
| actual HBD |
| error HBD |
| predicted TPSA |
| actual TPSA |
| error TPSA |
| predicted # rot. bonds |
| actual # rot. bonds |
| error # rot. Bonds |
| predicted # arom. rings |
| actual # arom. Rings |
| error # arom. Rings |
| predicted # aliph. rings |
| actual # aliph. Rings |
| error # aliph. Rings |
| predicted Fsp3 |
| actual Fsp3 |
| error Fsp3 |
| predicted QED |
| actual QED |
| error QED |
| predicted SA |
| actual SA |
| error SA |
